# Unraveling diversity by isolating peptide sequences specific to distinct taxonomic groups

**DOI:** 10.1101/2025.02.05.636664

**Authors:** Eleftherios Bochalis, Michail Patsakis, Nikol Chantzi, Ioannis Mouratidis, Dionysios Chartoumpekis, Ilias Georgakopoulos-Soares

**Affiliations:** Institute for Personalized Medicine, Department of Biochemistry and Molecular Biology, The Pennsylvania State University College of Medicine, Hershey, PA, USA; Department of Internal Medicine, Division of Endocrinology, Medical School, University of Patras, Patras, Greece; Huck Institute of the Life Sciences, Pennsylvania State University, University Park, PA, USA

## Abstract

The identification of succinct, universal fingerprints that enable the characterization of individual taxonomies can reveal insights into trait development and can have widespread applications in pathogen diagnostics, human healthcare, ecology and the characterization of biomes. Here, we investigated the existence of peptide k-mer sequences that are exclusively present in a specific taxonomy and absent in every other taxonomic level, termed taxonomic quasi-primes. By analyzing proteomes across 24,073 species, we identified quasi-prime peptides specific to superkingdoms, kingdoms, and phyla, uncovering their taxonomic distributions and functional relevance. These peptides exhibit remarkable sequence uniqueness at six- and seven-amino-acid lengths, offering insights into evolutionary divergence and lineage-specific adaptations. Moreover, we show that human quasi-prime loci are more prone to harboring pathogenic variants, underscoring their functional significance. This study introduces taxonomic quasi-primes and offers insights into their contributions to proteomic diversity, evolutionary pathways, and functional adaptations across the tree of life, while emphasizing their potential impact on human health and disease.

## Introduction

The number of available reference proteomes has rapidly increased in recent years, a trend that is expected to continue in the foreseeable future (UniProt Consortium 2023). The availability of a large and diverse set of proteomes of different organisms provides an opportunity to increase our understanding of protein sequence and functional diversity in nature across taxonomic groups. Such research could reveal insights in trait development, through findings pertaining to sequence conservation and divergence mechanisms, the emergence of proteins with new functional roles and can have applications in biomarker discovery, pathogen surveillance and human health among others (Al-Amrani et al. 2021; Lacerda and Reardon 2009). Such advances can be facilitated if the availability of ever-expanding proteomic information is coupled with novel and insightful algorithms to process this abundance of biological information.

Peptide k-mers are defined as oligopeptide sequences of length *k* and are often used in proteomics analyses (Moeckel et al. 2024). The number of possible peptide k-mers exponentiates with k-mer length, leading to oligopeptide sequence uniqueness even at low k-mer lengths (Mouratidis et al. 2024; Georgakopoulos-Soares et al. 2021a). Because of their ease of identification, peptide k-mers have been implemented in a number of applications including in mass-spectrometry-based proteomics (Chapman 2013), motif search and evolutionary studies (Wen et al. 2014), for taxonomic classification, antimicrobial resistance and pathogen detection (ValizadehAslani et al. 2020) and for the identification of therapeutic targets (Wu et al. 2019; Hajisharifi et al. 2014) among others.

Nullpeptides are the shortest oligopeptide sequences absent from a proteome (Hampikian and Andersen 2007). We and others have previously provided evidence that there are selection constraints against certain nullpeptides (Georgakopoulos-Soares et al. 2021a; Koulouras and Frith 2021; Navon et al. 2016; Poznański et al. 2018). Nullpeptides have been previously used as cancer killing compounds (Alileche et al. 2012; Alileche and Hampikian 2017; Ali et al. 2024), indicating that they can be used as potential drugs, and a subset of them has been shown to recurrently emerge during cancer development, while neoantigens with nullpeptides have been shown to be more immunogenic (Tsiatsianis et al. 2024). Additionally, peptide primes are the subset of nullpeptide sequences that are absent from every proteome (Hampikian and Andersen 2007). Previous research has provided evidence that peptide primes are immunomodulators and can enhance antigen specific immune responses in vaccine adjuvants (Patel et al. 2012). These studies exemplify the utility of different sets of k-mer peptides across biological problems.

We recently described the concept of quasi-prime peptides, which are the shortest peptides that are unique to a species’s proteome and are absent from every other proteome known (Mouratidis et al. 2023). We demonstrated that quasi-prime peptides can be identified across taxonomic groups at six and seven amino acid kmer lengths and characterize the set of proteins that harbor them. Here, we have extended the concepts of quasi-prime peptides to incorporate taxonomic groups, identifying k-mer peptides that are present in one or multiple species of a taxonomic group, but absent from every known proteome outside that group. Using 24,073 reference proteomes, we provide proof of the existence of k-mer peptides with this property at the superkingdom, kingdom and phylum levels **(Figure 1)**. We demonstrate the role of quasi-prime peptides in evolutionary divergence and taxonomic adaptations and provide evidence that human quasi-prime loci are more likely to harbor pathogenic variants, emphasizing their functional importance. Our study introduces taxonomic quasi-primes and provides evidence for their roles in shaping proteomic diversity, evolutionary trajectories, and functional adaptations across the tree of life, while underscoring their potential relevance to human health and disease.

**Figure 1:**
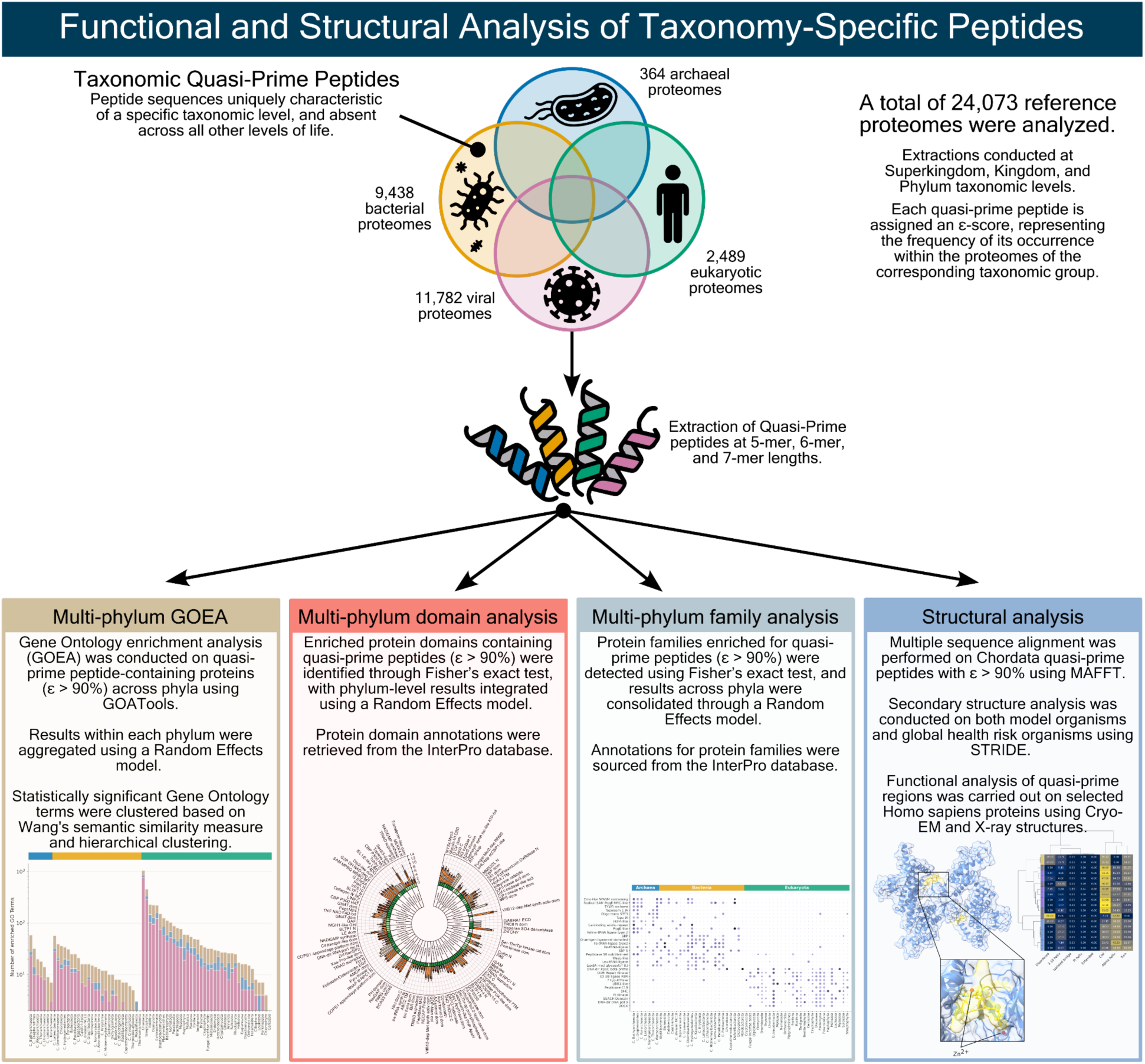
Overview of taxonomic quasi-prime peptide identification and analysis pipeline. Identification of taxonomic quasi-prime peptides was performed for 24,073 reference proteomes at the superkingdom, kingdom and phylum levels. To understand the roles of taxonomic quasi-prime peptides a thorough characterization of was performed, including gene ontology enrichment analyses, protein domain and family analyses and structural examinations.

## Materials and methods

### Proteomic datasets

Reference proteomes were obtained from UniProt (Release 2024_01), comprising a total of 24,073 species, including 364 archaeal, 9,438 bacterial, 2,489 eukaryotic, and 11,782 viral proteomes.

Peptide k-mer extraction was performed as previously described in (Mouratidis et al. 2023), for k-mer lengths of five to seven amino acids. We defined *T* as the superset of all considered taxonomies, *K* as a given k-mer and *P* as a proteome.

**Definition of k-mer.** We say that a k-mer *K* belongs to a taxonomic group *T*_*i*_ if and only if there exists at least one species *S* in *S* such that *T*_*i*_ is found in *S*, that is:

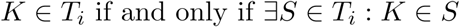

The set of all taxonomic quasi-primes of group *T*_*i*_ is then defined as:

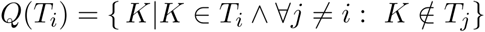

**Definition of Taxonomy.** For species classification, a taxonomy organizes species into hierarchical categories such as superkingdoms, kingdoms, and phyla.

#### Definition of ε-score

The ε-score for a k-mer *K* in a taxonomic group *T*_*i*_ is a measure of the frequency with which *K* appears in the proteomes of taxonomic group *T*_*i*_.

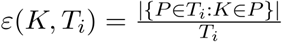

An ε-score equal to zero indicates complete absence of the k-mer *K* across all species of the taxonomic group *T*_*i*_, while an ε-score of one hundred indicates universal presence of k-mer *K* across all member species of the taxonomic group *T*_*i*_.

#### Definition of taxonomic quasi-primes peptides

Taxonomic quasi-prime peptides were defined as peptide sequences present in species of a taxonomic group and absent from all species outside that taxonomic group.

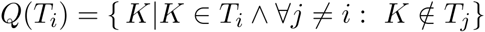

### Quasi-prime peptide protein matching

The Peptide Match command line tool (Chen et al. 2013) was used to map quasi-prime peptide sequences to the protein sequences containing them. The Lucene index needed for Peptide Match was created using UniProt Reference Proteome sequences containing SwissProt and TrEMBL entries. After the mapping process, a file containing the quasi-prime peptides of interest, the corresponding UniProt Accession ID, and the range containing the peptide was obtained.

### Species clustering based on Phylum ε-scores

Uniform Manifold Approximation and Projection (UMAP; McInnes, Healy, and Melville 2018) was employed using the Python library umap-learn (version 0.5.7) to analyze phylum quasi-prime 7-mers corresponding to the top 50th percentile of ε-scores within each phylum. This approach aimed to visualize how species clustered in two-dimensional space based on their taxonomic quasi-prime composition. A semi-supervised, density-based UMAP was implemented, incorporating a target weight of 0.25 for clustering based on labels. The algorithm parameters were configured with 30 neighbors and a minimum distance of 0.1 to optimize cluster resolution.

### Multi-species Gene Ontology Enrichment Analysis

Multi-species Gene Ontology Enrichment Analysis (MGOEA) was performed on quasi-prime peptide-containing proteins using GOATools v1.4.12 (Klopfenstein et al. 2018) at the Phylum level. A single Gene Ontology Enrichment Analysis (Ashburner et al. 2000) was performed for each Phylum species and results were combined using appropriate statistical methods (See Random effects model). The study population for each analysis consisted of the proteins with quasi-prime peptides, whereas the background population was represented by all the proteins expressed from the corresponding species. The Open Biological and Biomedical Ontologies (OBO) 1.4 file (.obo), containing ontology information and needed for the GOATools package, was obtained from the Gene Ontology Resource (Gene Ontology Consortium et al. 2023) (https://geneontology.org/, release 2024-09-08) and included a total of 40,939 GO terms and 7,894,411 annotations for 5,426 species. Finally, the Gene Ontology Annotation file (GAF) representing the relationship between UniProt Accession and Gene Ontology terms was downloaded from the Gene Ontology Annotation (GOA) Database version 222, released on 05 August, 2024 (https://ftp.ebi.ac.uk/pub/databases/GO/goa/UNIPROT/goa_uniprot_all.gaf.gz). Pre-MGOEA species were filtered to keep only those that possessed a protein count greater than 10, while also ensuring that protein counts were not propagated during the analysis. Post analysis only Gene Ontology terms with a p-value less than 0.05 were selected.

### Gene Ontology result clustering

Post-MGOEA enriched terms with an adjusted p-value less than 0.05 were clustered into broader representative GO terms to minimize redundancy and noise and highlight broader functional themes present across the phyla. Each GO term class (Biological Process, Molecular Function and Cellular Component) within each phylum was handled separately and Wang’s semantic similarity measure (Wang et al. 2007) was used to calculate the pairwise similarity of terms. These similarity values were converted to a distance matrix for agglomerative hierarchical clustering with the average linkage method. The optimal clustering threshold (Bettembourg, Diot, and Dameron 2015) specifically for each GO term class was set to 0.54 for biological processes, 0.535 for molecular functions and 0.52 for cellular components. Post-clustering a representative term for each cluster was selected based on the adjusted p-value and ties between terms were resolved using the presence percentage across phylum-specific species and the combined LOR, where terms with the highest value were characterized as representative. For the representative terms a weighted average LOR was calculated using the standard error as weight followed by winsorization at the 95th percentile to minimize the effect of extreme values. GO terms that were not assigned to any cluster were also retained, if the term’s adjusted p-value was less than 0.05.

### Multi-species Protein Entry Enrichment Analysis

Quasi-prime peptide-containing proteins were subjected to multi-species Protein Entry Enrichment Analysis (MPEA), where the presence of quasi-prime peptides within functional protein domains as well as the protein family composition of these proteins was analyzed. Protein domain and protein family data were obtained from the InterPro database (version 101.0 updated on 25th July 2024) (Paysan-Lafosse et al. 2023), which contains 14,950 domains and 26,089 family entries. For the enrichment analysis, Fisher’s exact test was implemented, from which Odds Ratio (OR) and a p-value was calculated for each entry, which were downstream combined to a final value that represents the effect size across Phylum species. Haldane-Anscombe correction (Agresti 1999) was applied to all cells of the 2×2 contingency table used in Fisher’s exact test to account for 0 values, which may lead to infinite estimates. Only entries with a p-value less than 0.05 OR greater than 1 and with presence across more than 1 species were subjected to effect combination, whereas entries present only in one species were retained, if they passed the p-value and OR cut-off.

### Enrichment combination across species using Random Effects Meta-Analysis method

A meta-analysis technique employing a random effects model was performed to identify gene ontology terms and functional entries (domains or families) that are present across multiple species of the same Phylum and evaluate the combined enrichment of each of them. Common items were filtered based on the criteria described earlier, and the natural logarithm of the OR value (LOR) was calculated. An original fixed-effect weight was calculated for each study (species), so that studies with more precise estimates (smaller standard error) are given a larger weight in the analysis, as follows:

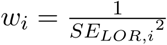

where, *w*_*i*_ represents the weight for the i_th_ study, *SE*_*LOR,i*_ is the standard error of the LOR value for the ith study and *SE*_*LOR,i*_ is the variance of the LOR estimate for the i_th_ study. The initial combined enrichment was obtained through the computation of the fixed effects weighted mean, which will be later used to assess the heterogeneity between-studies. The calculation of the fixed effects weighted mean goes as follows:

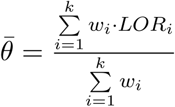

where *θ* represents the weighted mean LOR using the fixed-effects model and *k* represents the total number of studies. The use of Cochran’s Q statistic was implemented to measure the total variability in enrichment values across studies and will be used to to estimate variance due to heterogeneity. This formula was used to calculate the Q-statistic:

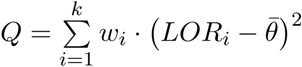

where *Q* is Cochran’s Q statistic and the term 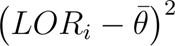 represents the squared deviation of each study’s effect size from the weighted mean. A constant was computed to adjust the variance of the calculated weights and also for the estimation of the between-study variance. The calculation goes as follows:

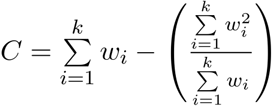

where *C* is the constant,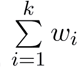 is the sum of weights obtained from all studies and 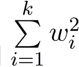 is the sum of the squared weights. A fraction adjustment is performed, because it accounts for the variability of the weights. As implied earlier, between study-variance was estimated using the Tau-squared statistic, due to its ability to calculate the amount of variance in enrichment values due to real differences between studies rather than chance and it can ensure that the variance estimate is non-negative. The t^2^ statistic is calculated using this formula:

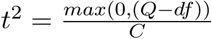

where *t*^2^ is the heterogeneity variance and *df* represents the degrees of freedom (k-1) used to assess the statistical significance of the *Q* statistic against the chi-squared distribution. The final weights used for the combined enrichment values were obtained using the following random effects model:

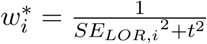

*w*^*^_*i*_ is the final adjusted weight for the i^th^ study using the random effects model and *SE*_LOR,i_ is the within-study variance of the i^th^ study. The computation of the combined overall enrichment value for each item using the random effects model is represented below:

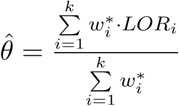

*θ* is the combined enrichment value (LOR) using the random effects model, the numerator represents the sum of adjusted LOR values while the denominator represents the sum of the adjusted weights. In all these calculations, we have assumed that taking the natural logarithm of the odds ratio provides a consistent measure of enrichment effect size. This assumption enables us to extend the enrichment analysis across species and calculate the final combined enrichment value for each item. The final step to this random-effects model was to calculate the statistical significance of the combined enrichment value against the null-hypothesis that the combined enrichment value is zero. To achieve this, first we calculated the standard error of the combined enrichment value, then we computed a modified Z-score, and with this Z-score a final p-value was obtained. The formula used for modified Z-score is the following:

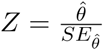

where a larger absolute value of *Z* indicates a more significant deviation from the null-hypothesis and the formula for the p-value calculation:

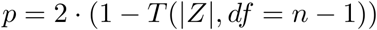

where *P* represents the calculated p-value and *T*(|*Z*|) denotes the cumulative distribution function of the t-distribution with n-1 degrees of freedom. The choice to calculate p-values based on the t-distribution was made to address the small sample size of certain underrepresented phyla. Items were filtered to keep only those with a meta-analysis adjusted p-value less than 0.05. Multiple testing correction was applied using the Benjamini-Hochberg procedure.

### Secondary and tertiary structure analysis

Quasi-prime peptides were subjected to structural analysis. Original PDB files of the proteins containing quasi-prime peptides were obtained from the AlphaFold Protein Structure Database (Jumper et al. 2021; Varadi et al. 2024) updated as of September 2024. Quasi-prime peptide regions from each protein were extracted and their secondary structure was identified using the STRIDE (STRuctural IDEntification) algorithm (Frishman and Argos 1995) as provided by the ssbio v0.9.9 tool package. Quasi-prime peptides that presented with no hydrogen-bonds were characterized to have a Disordered conformation. PDB files presenting the interaction between protein and ligands were downloaded from the RCSB Protein Data Bank (Berman et al. 2000) (updated as of 2024, download timestamp: November 21st 2024), containing a total of 225,681 available structures. Only conformations obtained through Cryo-EM or X-Ray were selected. Protein visualization and hydrogen-bond detection between ligand and protein was performed with the use of the UCSF ChimeraX version 1.8 software (Meng et al. 2023).

#### Protein ortholog mapping and Multiple Sequence Alignment

Protein orthologs for Homo Sapiens proteins across the Chordata phylum were obtained through the EggNOG v5.0.0 (Huerta-Cepas et al. 2019) database, which contains 4.4 million orthologous groups and data for 5,090 Organisms and 2,052 Viruses. The orthologous groups of choice were later subjected to Multiple Sequence Alignment (MSA) using the MAFFT version 7 command line tool (Katoh and Standley 2013) with the options: --localpair --maxiterate 1000 --amino --thread 5. The resulting alignments were trimmed using ClipKIT v.2.3.0 (Steenwyk et al. 2020) with the options -smart-gap to remove poorly aligned protein regions and improve the phylogenetic signal by focusing on well-conserved segments. The trimmed results were visualized using Jalview v2.11.4.1 (Waterhouse et al. 2009). Sequences were ordered based on pairwise similarity, quasi-prime-containing regions were highlighted whereas distant areas that did not contain quasi-prime peptides were hidden from the visualization.

#### Pathogenicity prediction in taxonomic quasi-prime protein regions

To predict the pathogenicity of single-nucleotide missense variants, we employed AlphaMissense (J. Cheng et al. 2023), (version v3, updated as of September 19, 2023, https://zenodo.org/records/10813168). The analysis included all possible missense variants (approximately 71 million) derived from 19,000 canonical protein-coding transcripts in the human genome (hg38 build). Our study specifically focused on human proteins containing taxonomic quasi-prime 7-mers with ε-scores exceeding 90%. We conducted a comparative analysis of the pathogenicity associated with missense mutations located within these taxonomic quasi-prime loci and those occurring outside these loci. Mutations with AlphaMissense scores below 0.1 were classified as likely benign, while those with scores above 0.9 were designated as highly pathogenic.

## Results

### Identification of sequences that are unique to individual taxa

It remains unknown if peptide k-mer sequences which are unique to a particular taxonomic group play a role in the emergence of novel traits within individual taxa. Here, we investigated the potential presence of taxonomic quasi-prime peptide sequences, which are k-mer peptide sequences that are found in proteomes of a single taxon, and otherwise absent from all other taxa. We performed this investigation at the superkingdom, kingdom and phylum levels for k-mer lengths of five, six and seven amino acids in all organisms with an available reference proteome, totalling 24,073 reference proteomes spanning the tree of life. K-mer lengths below five amino acids were not considered, as tetrapeptides are highly prevalent and in the human proteome all possible tetrapeptides are observed (Georgakopoulos-Soares et al. 2021b; Chantzi et al. 2024). Octapeptides and longer oligopeptides were not considered as the possible proteome space becomes extremely large (20^8^), limiting the set of k-mers that are shared between multiple species in a taxonomy.

### Derivation of superkingdom- and kingdom- specific quasi-prime peptides

First, we define taxonomic quasi-primes, sequences specific to a taxonomy and define the ε-score, representing the percentage of species of the taxonomy in which a peptide k-mer is found in. (see Methods). Taxonomic quasi-prime five-mers are identifiable at the superkingdom level and are exclusively found within the Eukaryota. We identified 12 distinct five-mers with ε-scores ranging from 0.52% to 3.74%, and a median ε-score (ε_M_) of 1.67% **(Supplementary** Figure 1). The sequences of these 12 peptides are detailed in **Supplementary Table 1**. Regarding taxonomic quasi-prime six-mers, we find that the median percentage of species containing these six amino acid peptides is 0.01% in viruses, 0.02% in bacteria, 0.24% in eukaryotes, and 0.27% in archaea (**Figure 2a**.) Similar patterns are observed for seven-amino acid taxonomic quasi-prime peptides (**Figure 2a**). The larger number of taxonomic quasi-primes identified in Eukaryotes stems from their larger proteome sizes (Spearman’s correlation coefficient ρ=0.963, p-value<0.001) **(Supplementary** Figure 2). Additionally, due to the large k-mer space these findings translate to a considerable number of superkingdom-specific peptides. Specifically, at six and seven amino acids peptide k-mer length, we observe thousands of peptides that are only found in species of a single superkingdom or only in Viruses (**Figure 2a**). For instance, we observe 85,373 and 86,483,511 peptides found uniquely in bacterial proteomes at six and seven amino acids k-mer lengths respectively (**Figure 2a**).

**Figure 2:**
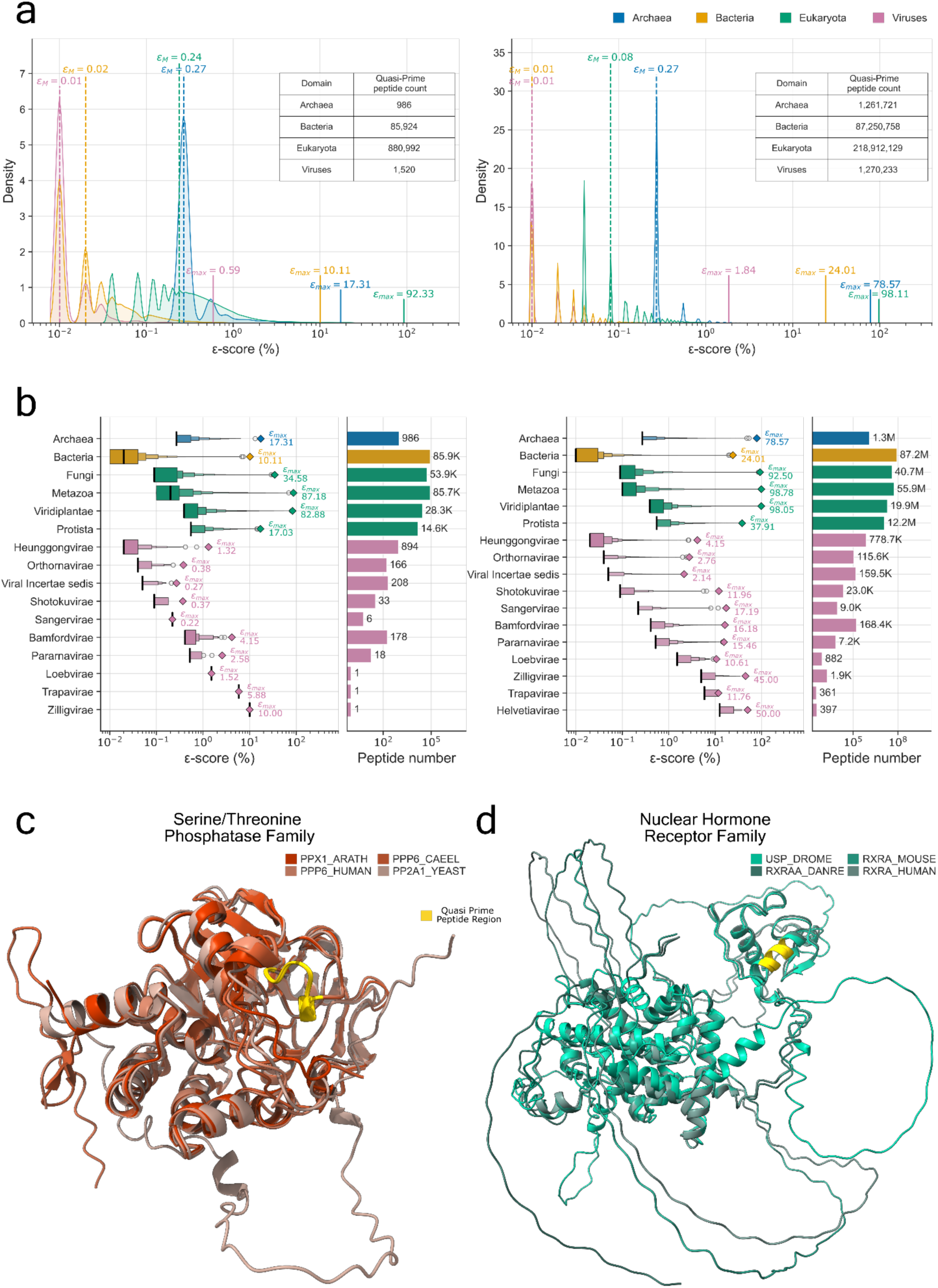
Quasi-prime peptide count and ε-score distribution. **a** Kernel Density Estimate plots showing quasi-prime peptide ε-score distribution at Superkingdom level. Dotted lines represent the median ε-score (ε_M_) and solid lines represent the maximum ε-score (ε_max_) value. Tables inside each plot display the number of unique quasi-prime peptide counts. **b** Quasi-prime peptide ε-score distribution and peptide count at Kingdom level. Left to right: Letter-value plot of ε-score distribution, where the ε_M_ value is depicted as a solid black line and the ε_max_ value as a rhombus. Barplot of the unique quasi-prime peptide counts. (Data for quasi-prime peptide 6mers are displayed to the left and quasi-prime peptide 7mers are displayed to the right). **c** Superposition of proteins representing the Serine/Threonine Phosphatase family at the eukaryotic superkingdom. Results are shown for *Arabidopsis thaliana, Caenorhabditis elegans, Homo sapiens*, and *Saccharomyces cerevisiae* orthologs. **d** Superposition of proteins representing the Nuclear Hormone Receptor family at the metazoan kingdom. Results are shown for *Drosophila melanogaster*, *Mus musculus*, *Danio rerio*, and *Homo sapiens* orthologs. Taxonomic quasi-prime peptides are marked in yellow.

We also find that the six amino acid taxonomic quasi-prime peptides found in the largest number of species (ε_max_) in each of the three superkingdoms and in viruses were in 0.59%, 10.10%, 17.31% and 92.33% of the species in viruses, bacteria, archaea and eukaryotes respectively. This means that we can identify one six-mer peptide found in 92.33% of eukaryotes and otherwise absent from all bacteria, archaea and viruses with reference proteomes available. Interestingly, for peptide lengths of seven amino acids we are able to identify ε_max_ of 98.11% for eukaryotic species for the peptide sequence SAPNYCY, which maps to proteins belonging to the serine/threonine phosphatase family, and are highly conserved in eukaryotes (Ohama 2019) (**Figure 2a,c**). The observed patterns underscore the significance of taxonomic quasi-prime peptides in distinguishing between superkingdoms, offering valuable insights into proteomic diversity across different domains of life and holding potential as molecular markers for superkingdom-level classification.

Next, we analyzed the distribution of taxonomic quasi-primes across various organismal and viral kingdoms, observing significant variability within individual groups. This variability was particularly pronounced in viral kingdoms, where ε_M_ values ranged from 0.02% to 10.00% at the six-mer level and from 0.02% to 5.00% at the seven-mer level in Heunggongvirae and Zilligvirae, respectively (**Figure 2b**). Among eukaryotic kingdoms, the lowest ε_M_ value was found in Fungi (0.09%), while the highest was observed in Protista (0.55%) **(Figure 2b)**. Further analysis of ε_max_ at the six-mer level within these kingdoms revealed values ranging from 17.03% in Protista to 87.18% in Metazoa. For seven-mers, ε_max_ values reached up to 37.91% in Protista and 98.78% in Metazoa **(Supplementary Table 1)**. We identified the taxonomic quasi-prime seven-mer, CKGFFKR, which represents the ε_max_ within the Metazoan kingdom. This seven-mer was mapped to proteins associated with the nuclear hormone receptor family, whose genes show strong sequence conservation and little evidence for positive selection in Metazoans (Krasowski et al. 2005) (**Figure 2b,d**).

We conclude that we can identify highly superkindom-specific and kingdom-specific peptides shared in a significant proportion of the species of those taxonomic groups. We note, however, that the number of taxonomic quasi-prime peptides we can detect is influenced by the proteome size and the number of species available within each superkingdom and kingdom. As more reference proteomes become available, this representation will become increasingly complete.

### Derivation of phylum-specific quasi-prime peptides

We investigated the presence and distribution of taxonomic quasi-primes across various phyla. At the six-amino-acid sequence length, ε_M_ values displayed substantial variation. In eukaryotic phyla, values ranged from 0.14% in Ascomycota species to 100.00% in Foraminifera and sixteen other eukaryotic phyla. For bacterial phyla, ε_M_ values spanned from 0.03% in Pseudomonadota to 100.00% in Abditibacteriota and 17 additional bacterial phyla. Archaeal phyla exhibited values between 0.43% in Euryarchaeota and 100.00% in seven candidate archaeal phyla. In viral phyla, ε_M_ values ranged from 0.02% in Uroviricota to 10.00% in Taleaviricota (Figure 3**-4; Supplementary** Figure 3). A similar distribution pattern was observed for the seven-amino-acid sequence length. In eukaryotic phyla, Ascomycota again exhibited the lowest ε_M_ value at 0.14%, while Porifera reached the maximum value of 100.00%, shared with 16 other eukaryotic phyla. Among bacterial phyla, ε_M_ values ranged from 0.03% for Pseudomonadota to 100.00% for Abditibacteriota and 18 additional bacterial phyla. For archaeal phyla, Euryarchaeota showed the lowest ε_M_ value of 0.43%, while Candidatus Bathyarchaeota and seven other candidate archaeal phyla attained ε_M_ of 100.00%. Viral phyla exhibited values ranging from 0.02% in Uroviricota to 12.50% in Dividoviricota (Figure 3**-4; Supplementary** Figure 3). We also observe that superkingdoms cluster by taxonomic quasi-primes (Figure 3b). The complete distribution of ε_M_ values alongside ε_max_ for each phylum is available at the **Supplementary Table 3**.

**Figure 3:**
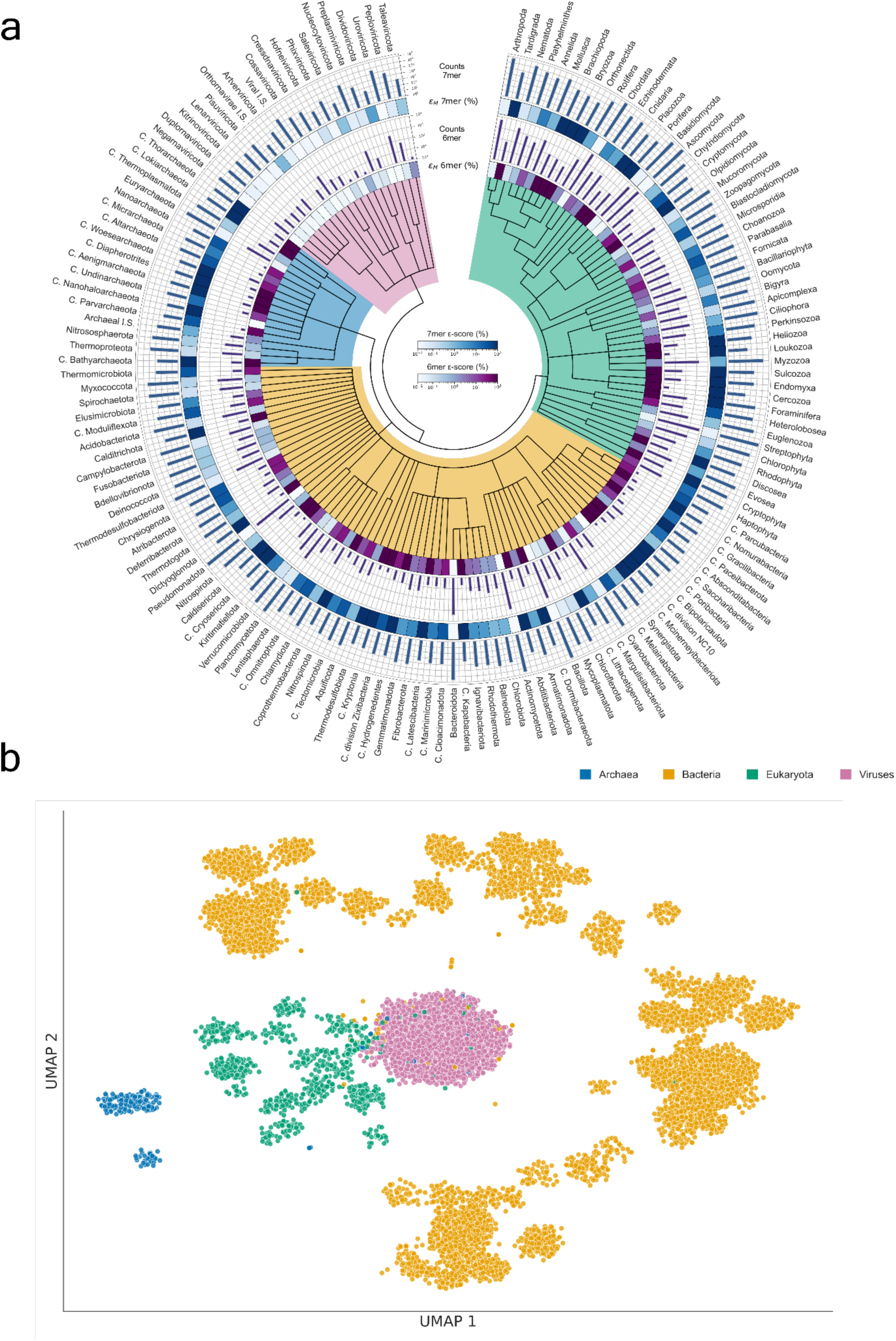
Quasi-prime peptide statistics at Phylum level. **a** Circos plot depicting the phylogenetic organization of Superkingdoms, Kingdoms and Phyla at the Archaeal, Bacterial, Eukaryotic and Viral levels (center). The number of unique quasi-prime peptides, alongside the median ε-score value (εM) at 6mer and 7mer length is also shown. Inner to outer: εM for 6mer quasi-prime peptides; total 6mer quasi-prime peptide counts; εM for 7mer quasi-prime peptides; total 7mer quasi-prime peptide counts. b UMAP plot depicting the clustering of reference proteomes based on their taxonomic quasi-prime 7mers. Only the top 50th percentile of quasi-primes based on their ε-score was used for the clustering.

**Figure 4:**
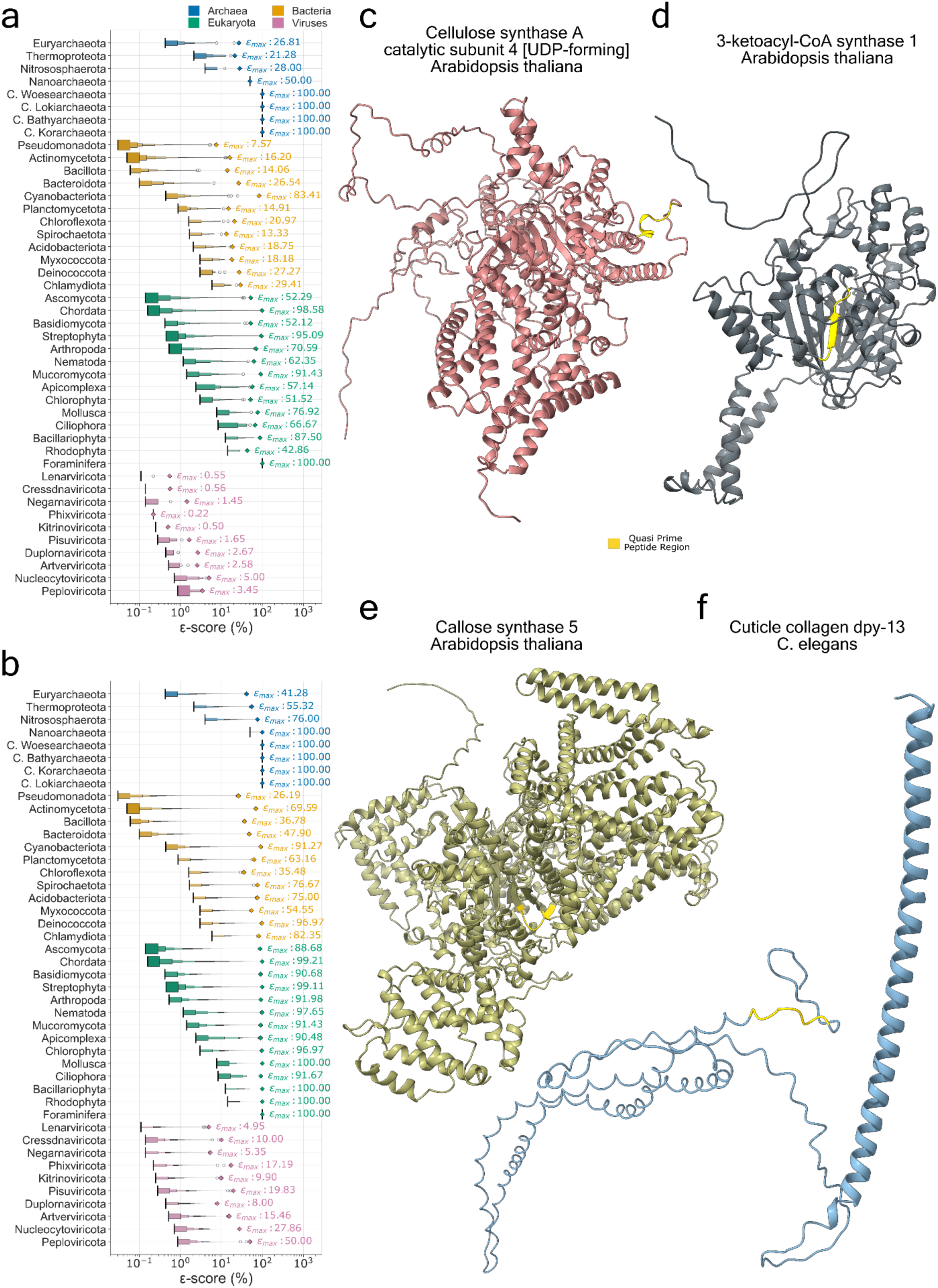
ε-score distribution of quasi-prime peptides across representative phyla at the superkingdom level. Letter-value plots illustrate the ε-score distributions for representative phyla at the superkingdom level. The εM value is depicted as a solid black line and the εmax value as a rhombus. Distributions are shown separately for taxonomic quasi-prime peptides of different lengths: a Taxonomic quasi-prime six-mers b Taxonomic quasi-prime seven-mers. The color represents the corresponding superkingdom. c-f Protein structure of: c the Cellulose Synthase A Catalytic Subunit 4, d 3-Ketoacyl-CoA Synthase 1, e Callose synthase 5, in Arabidopsis thaliana and cuticle collagen DPY-13 protein of C. elegans. Taxonomic quasi-prime peptides are marked in yellow.

Across all three cellular superkingdoms, several phyla exhibited an ε_max_ of 100.00%, whereas for viral phyla the ε_max_ was capped at 50%. This is likely the result of viruses’ rapid evolution in response to host immune pressures resulting in viral phyla encompassing greater genetic variation than cellular phyla. For both Streptophyta and Nematoda, we analyzed the proteins from which these highly phylum-specific peptide seven-mers originated. In Streptophyta, three taxonomic quasi-primes were identified with an ε_max_ of 99.11%. These sequences, TPWPGNN, EHFCIHA, and THHEYIQ, were found in *Arabidopsis thaliana* within the proteins Cellulose Synthase A Catalytic Subunit 4 (UDP-forming), 3-Ketoacyl-CoA Synthase 1, and Callose Synthase 5, respectively (Figure 4c-e). In Nematoda, a single peptide, ICPKYCA, was identified with an ε_max_ of 97.65%. This peptide is located in the cuticle collagen DPY-13 protein of *C. elegans* (Figure 4f), which is crucial for cuticle formation, serving both as an exoskeleton and a protective barrier against environmental challenges.

### Taxonomic quasi-primes enable the detection of loci divergence and functional adaptations across taxa

To explore the functional roles of taxonomic quasi-prime peptides, we conducted a GO enrichment analysis tailored to individual taxa. We developed an phylum-wide enrichment estimation via a Random Effects Meta-Analysis model (see Methods). The analysis focused on seven-mer quasi-primes with ε-scores above 90% for each phylum in Archaea, Bacteria, and Eukaryotes, emphasizing representative phylum-wide functional enrichments.

We identified statistically significant (adjusted p-value < 0.05) enriched GO terms across Biological Processes (BP), Molecular Functions (MF) and Cellular Components (CC) (Figure 5a). BPs were the most enriched GO class, followed by MFs, with CCs being less prevalent. Eukaryota, led by Chordata and Streptophyta, exhibited the highest number of enriched GO terms, particularly in BPs, while Archaea and Bacteria showed comparable numbers, reflecting their complexity, proteome size, and horizontal gene transfer (Figure 5a). Within Archaea, Candidatus Bathyarchaeota, one of the most prevalent microorganisms on Earth (Feng et al. 2019), exhibited a high number of enriched BPs (Figure 5a), likely due to its extensive metabolic versatility, linked to protein degradation, glycolysis, and the Wood–Ljungdahl pathway (Feng et al. 2019).

**Figure 5:**
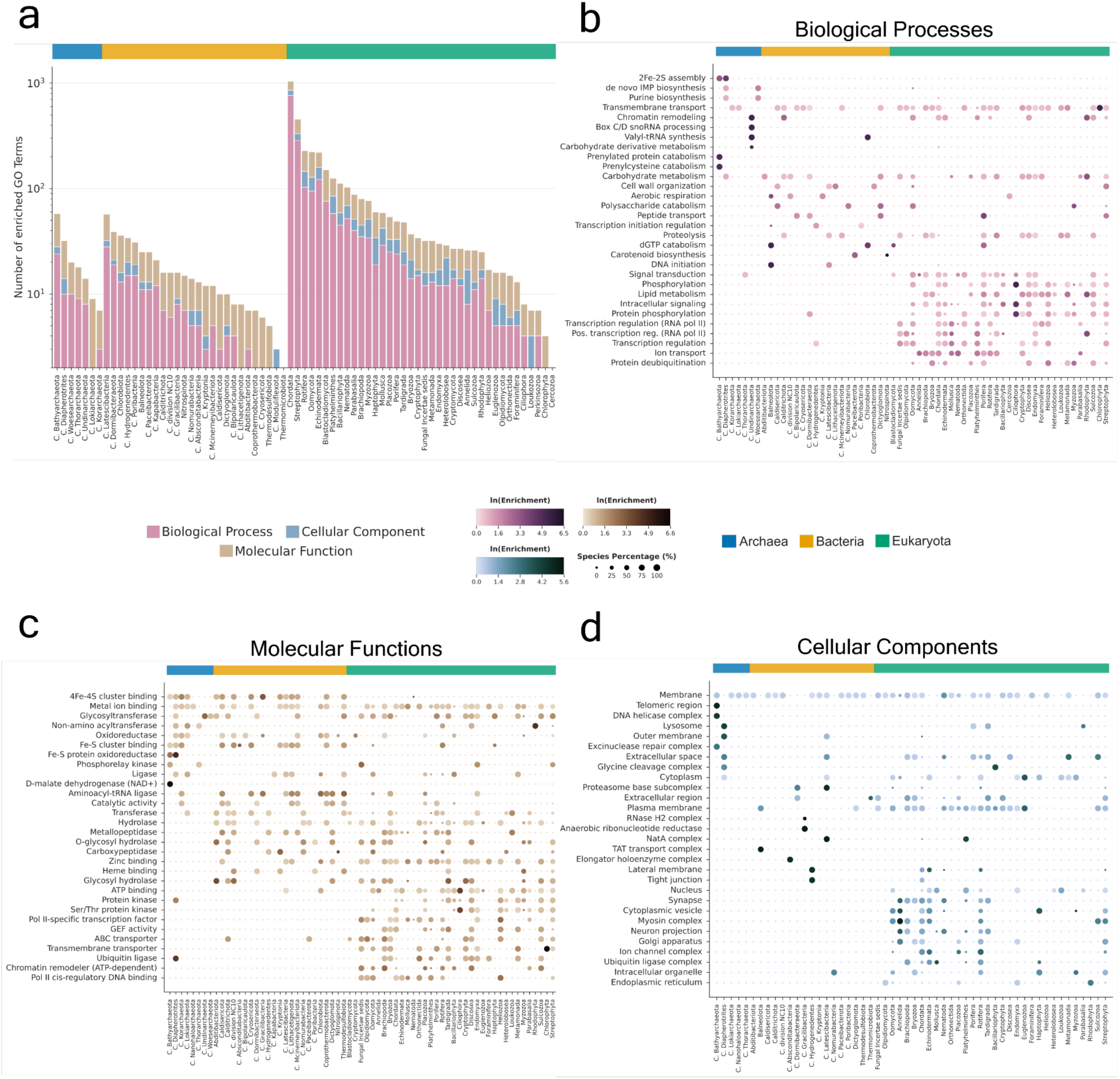
Multi-phylum Gene Ontology (GO) enrichment analysis at superkingdom level. **a** Stacked bar chart illustrating the total number of enriched GO terms, categorized by GO class, at cellular organism superkingdom level. b-d Heatmaps highlighting the top ten GO class specific terms most broadly enriched across phyla within each of the three cellular organism superkingdoms. Dot size indicates the prevalence of each GO term among species within a given phylum, while dot color denotes the combined natural log (ln) of enrichment values, reflecting the strength of enrichment. Heatmaps are organized as follows: b Biological Processes c Molecular Functions d Cellular Components.

We examined the prevalence of specific GO terms across multiple phyla within each superkingdom. Transmembrane transport was enriched across all three superkingdoms (Figure 5b), highlighting lineage-specific conservation. Eukaryotic-exclusive terms, such as RNA polymerase II transcription regulation, were absent in prokaryotes. Ion transport was enriched in eukaryotes with nervous systems, such as Chordata and Mollusca (Figure 5b). Key MFs, like metal ion binding and glycosyltransferase activity, possessed unique quasi-primes across superkingdoms, reflecting their role in fundamental cellular roles (Figure 5c). Examples of metal ion binding include oxidoreductase activity requiring iron ions in prokaryotic respiration and energy transduction (Barquera 2014), zinc ion binding for eukaryotic transcription factors (Kamaliyan and Clarke 2024), and glycosyltransferase activity for carbohydrate modifications using manganese ions (Breton et al. 2006) (Figure 5c). Enriched 4Fe-4S cluster binding in Archaea and Bacteria highlighted their role in anaerobic respiration and redox regulation (Beinert, Holm, and Münck 1997; Ibrahim et al. 2020). Membrane CC was enriched across all cellular superkingdoms showing showcasing adaptations in membrane protein structures through quasi-primes (Figure 5d).

To uncover shared functional themes, we analyzed top enriched GO terms for representative Phyla, which were clustered into broader categories using Wang’s semantic similarity measure and hierarchical clustering (Figure 6a-c). In Chordata, receptor tyrosine kinase signaling and ephrin receptor binding (Figure 6a-b) were enriched, crucial for nervous system development through axon guidance (Kullander et al. 2001; Kao and Kania 2011) and angiogenesis (N. Cheng, Brantley, and Chen 2002). Clathrin binding, alongside synaptic vesicles and postsynapse, were enriched, highlighting the role of clathrin-mediated endocytosis in Chordata (McMahon and Boucrot 2011) (Figure 6b-c).

**Figure 6:**
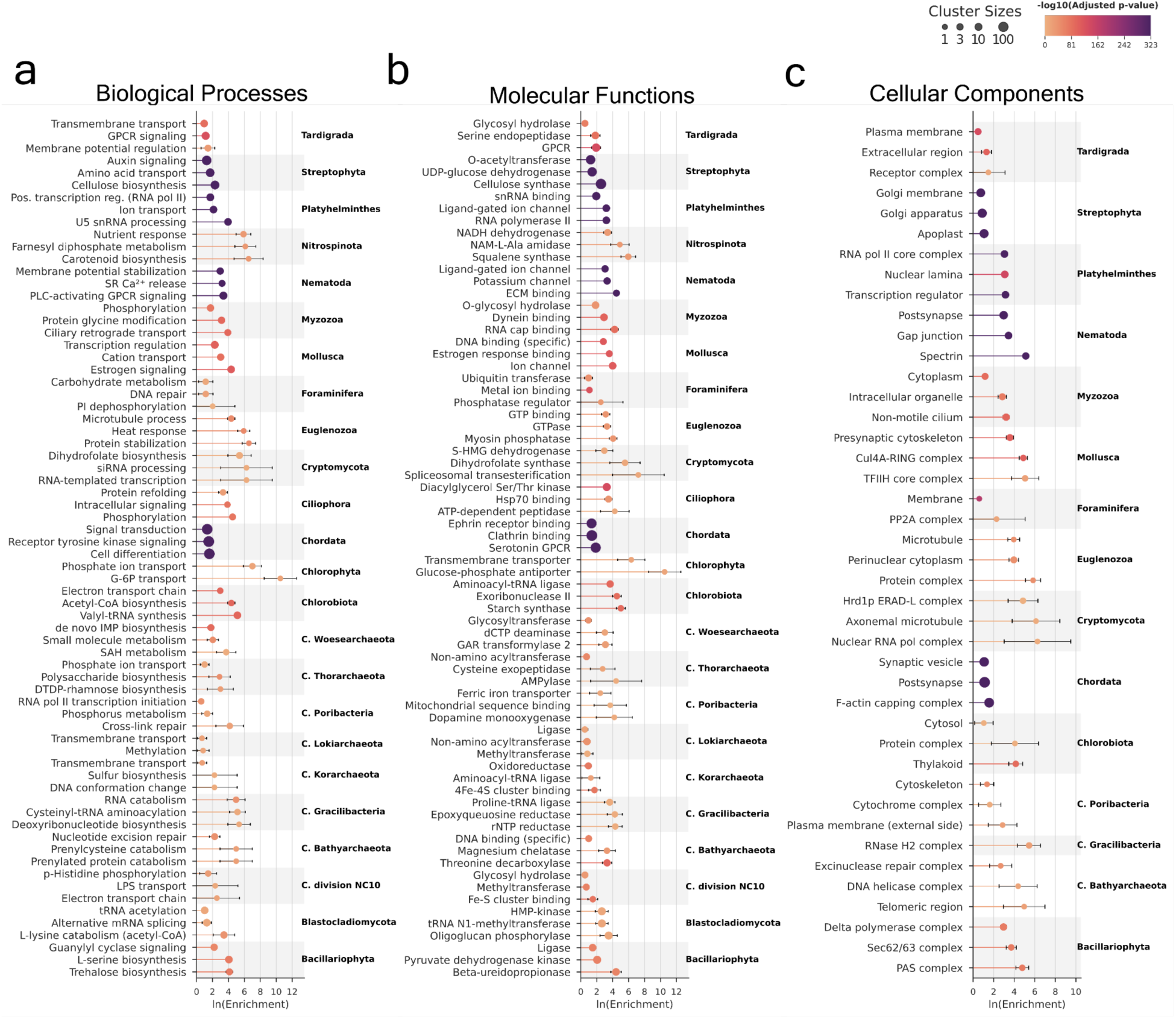
Gene Ontology (GO) Term Enrichment Analysis across superkingdom representative phyla, with taxonomic quasi-prime peptides. Lollipop plots display the mean enrichment of the top three most enriched clustered GO terms for representative phyla of the cellular organism superkingdoms, categorized into: a Biological Processes, b Molecular Functions, c Cellular Components. Only GO terms with an adjusted p-value less than 0.05 and a species representation greater than 5% are included. Dot size represents cluster size, lollipop color represents the −log10 adjusted p-value and error bars show the 95% confidence interval of the calculations. GO terms have been grouped into broader clusters using Wang’s semantic similarity measure combined with hierarchical clustering.

Mollusca showed enrichment of estrogen signaling, since they rely on environmental uptake of estrogen hormones (Balbi, Ciacci, and Canesi 2019). Poribacteria, a symbiotic phylum associated with Porifera, showed unexpected enrichment of GO terms related to RNA polymerase II transcription initiation BP and dopamine monooxygenase MF (Figure 6a-b). Poribacteria are known to mimic Porifera species and possess eukaryotic-like protein domains (Kamke et al. 2014), a property that appears to extend to taxonomic quasi-primes as well. In Nematoda, enriched GO terms included G-protein coupled receptor signaling (PLC-activating GPCR), linked to calcium release and muscle contraction, and postsynapse CCs (Liu et al. 2021) (Figure 6b-c).

Cytoskeletal GO term clusters were enriched with taxonomic quasi-primes across multiple phyla (Figure 6c). For instance, spectrin was enriched in Nematoda for stabilizing sensory neurons (Krieg, Dunn, and Goodman 2014), and the F-actin capping complex CC was critical for cytoskeletal remodeling in Chordata (Cooper and Sept 2008). Cytoskeletal components were enriched in other phyla, such as Euglenozoa (microtubules) and Cryptomycota (axonemal microtubules), essential for flagellar motility and host interactions (Jones et al. 2011).

These findings show the unique functional roles of taxonomic quasi-prime peptides with a high ε-score across diverse taxa, with enriched GO terms reflecting adaptations to specific cellular and environmental demands within each superkingdom.

### Taxonomic quasi-primes detect superkingdom- and phylum-level protein adaptations

To investigate taxon-specific functional changes, we analyzed the distribution of taxonomic quasi-primes within protein domains and families, collectively termed as entries. Across all three superkingdoms, the major facilitator superfamily domain (MFS_dom), critical for transmembrane transport (Pao, Paulsen, and Saier 1998), emerged as a highly enriched domain (Figure 7a), underscoring its pivotal role in transport processes essential for cellular survival and adaptation (Complete protein domain names can be found at: **Supplementary Table 3**). This aligns with previously found enriched transmembrane transport BP across all three superkingdoms Archaea and bacteria displayed similar enrichment patterns in entries, which were largely absent in eukaryota, and vice versa (Figure 7a, Figure 8a). This indicates evolutionary conservation of entries across these two superkingdoms, due to their shared environmental challenges, horizontal gene transfer events and adaptive responses. Enriched domains included those associated with oxidoreductase activity, transferase activity, catalytic activity, and metal ion binding. Particularly, the radical S-adenosyl-L-methionine (rSAM) domain and its associated families (e.g., Elp3/MiaA/NifB-like, PqqE-like) (Figure 7a, Figure 8a) were enriched, reflecting their roles in essential enzymatic functions. These families belong to the rSAM enzyme superfamily (Frey, Hegeman, and Ruzicka 2008) and involve 4Fe-4S cluster binding (4Fe4S_Fe-S-bd), critical for catalysis.

**Figure 7:**
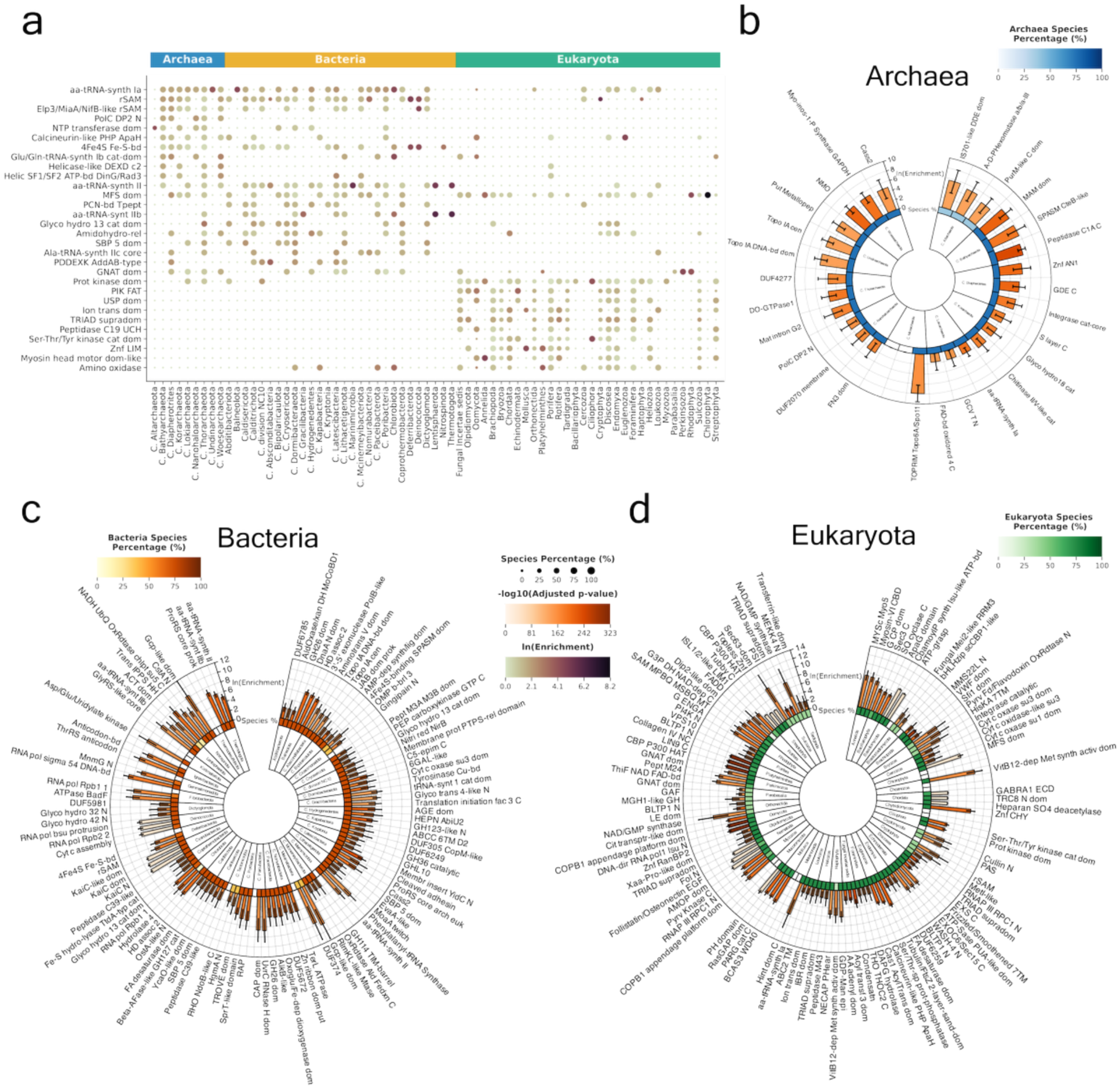
Preferential presence of taxonomic quasi-primes in specific protein domains. **a** Size- and color-coded heatmap displaying the top ten protein domains with taxonomic quasi-primes enriched across most phyla within each of the three cellular organism superkingdoms. Dot size indicates the prevalence of each protein domain within species of the respective phylum, while dot color corresponds to the combined ln(Enrichment) value for each domain. b-d Circos plots of the top three enriched protein domains per phylum with an adjusted p-value less than 0.05 and a species prevalence greater than 5% for each cellular organism superkingdom. b Archaea, c Bacteria, and d Eukaryotes. Inner to outer: Phylum name; Heatmap depicting the percentage of the species within the respective phylum that have the specific protein domain enriched; Barplot showing the ln(Enrichment) value with error bars representing the 95% Confidence Interval of the calculation.

**Figure 8:**
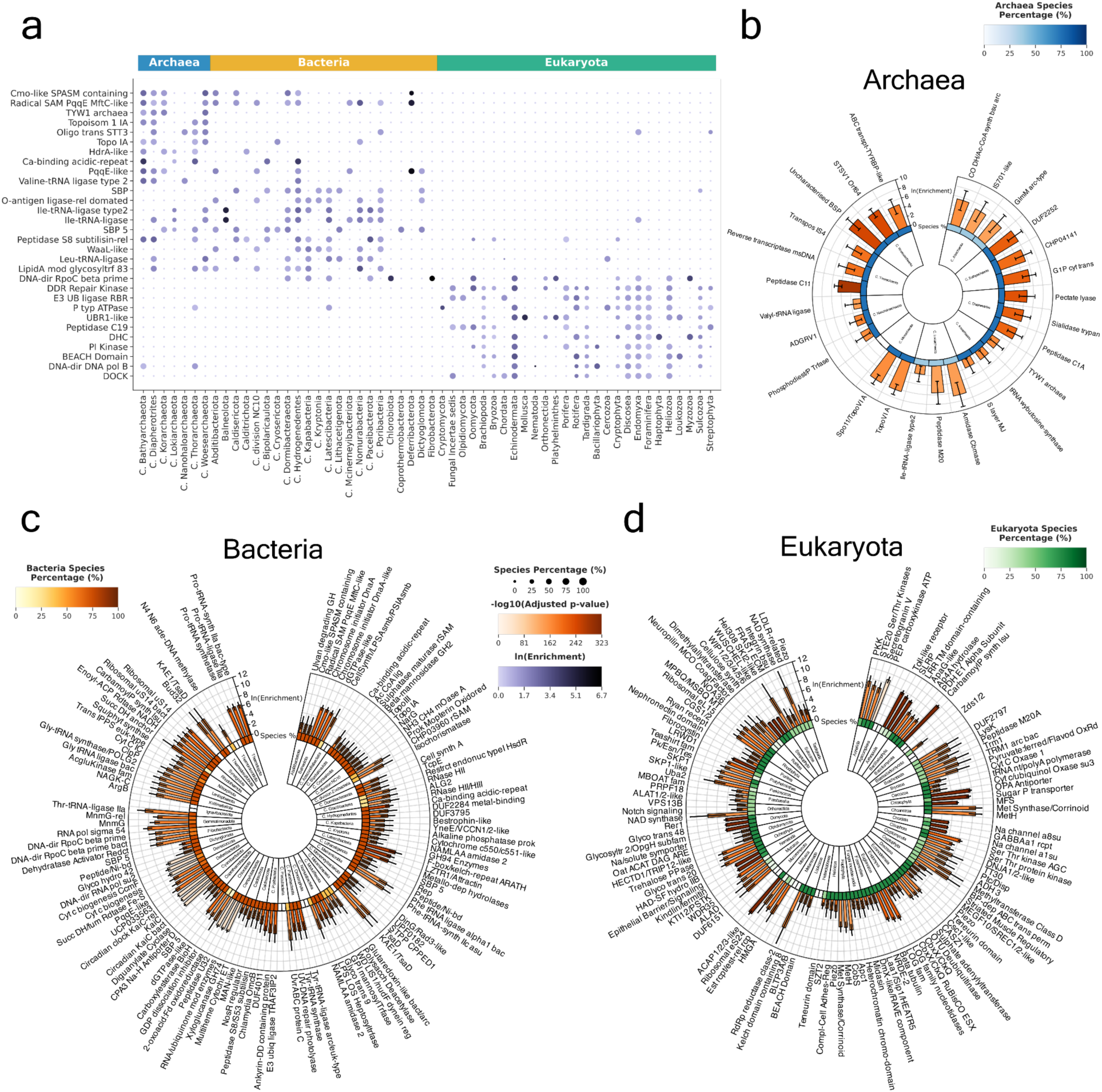
Analysis of the predominant taxonomic quasi-primes in superkingdom specific protein families. **a** Size- and color-coded heatmap identifying the top ten enriched protein families widely present across phyla within each of the three cellular organism superkingdoms. Dot size indicates the prevalence of each protein family within species of the respective phylum, while dot color corresponds to the combined ln(Enrichment) value for each family. **b-d** Circos plots showcasing the top three most enriched protein families per phylum with an adjusted p-value less than 0.05 and a species prevalence greater than 5% in: **b** Archaea, **c** Bacteria, and **d** Eukaryotes. Inner to outer: Phylum name; Heatmap depicting the percentage of the species within the respective phylum that have the specific protein family enriched; Barplot showing the ln(Enrichment) value with error bars representing the 95% Confidence Interval of the calculation.

Aminoacyl-tRNA synthetase domains were enriched in Archaea and Bacteria (aa-tRNA-synth Ia and aa-tRNA-synth II), with bacterial-specific enrichment of the class II G/P/S/T subtype (Figure 7a). Three class Ia tRNA ligase protein families (Valine-, Leucine and Isoleucine-tRNA ligases) are found enriched predominantly in most archaeal and bacterial phyla (Figure 8a), supporting protein synthesis under extreme conditions. These adaptations compensate for the limited diversity of post-transcriptional modifications in prokaryotes.

In Eukaryotes, enriched entries included the ion transport domain, (Figure 7a), found in sodium, potassium and calcium ion channels, the TRIAD supradomain present in the E3 ubiquitin ligase RBR family and the myosin head motor domain alongside the dynein heavy chain family (Figure 8a). E3 ubiquitin ligases are highly conserved across eukaryotes, since they are a part of the ubiquitin-proteasome system, involved in protein degradation and the maintenance of cellular homeostasis (Yang et al. 2021). Enrichment of motor protein entries highlights the necessity of quasi-prime adaptations for intracellular transport and cellular motility.

We also identified the top three enriched entries within specific phyla. In Cyanobacteria entries related to the circadian clock oscillator protein family (e.g. KaiC) (Figure 7b, Figure 8b) are enriched suggesting the high evolutionary conservation of quasi-primes in the regulation of day-night cycles (Markson and O’Shea 2009). In Candidatus Paceibacterota the UV-induced DNA damage repair photolyase family was enriched, warranting further study (Figure 8b). In Chordata, the Heparan sulphate-N-deacetylase protein domain, critical for the biosynthesis of heparan sulfate (Sarrazin, Lamanna, and Esko 2011), is highlighted (Figure 7c). Also, the sodium channel A8 and A1 subunit families, alongside the gamma-aminobutyric-acid A receptor, alpha 1 subunit are present across most chordata species further facilitating the involvement of quasi-primes in the function of the nervous system. Leucine-rich repeat families were prominent in Arthropoda (Figure 8c) underscoring roles in immune defense and development, with emphasis on Toll-like receptors (Dey et al. 2022) and extracellular matrix integrity (Matsushima, Miyashita, and Kretsinger 2021). Little is known about how these families function in Arthropods, and the existence of highly specific taxonomic quasi-primes emphasizes the need for further study.

### Secondary Structure Analysis of Taxonomic Quasi-Primes in Diverse Organisms

We next analyzed the secondary structures of protein loci containing taxonomic quasi-prime seven-mers with an ε-score greater than 90% across thirteen model organisms and seventeen pathogens of global health concern. Our findings revealed a distinct enrichment of taxonomic quasi-primes in protein coils, followed by alpha helices and turns (Figure 9a-b). We evaluated the statistical significance of this structural preference using Kruskal-Wallis tests. For coiled-like conformations (encompassing alpha helices, coils, and turns), the analysis yielded an H-statistic of 213,788.82 with a p-value < 0.001 in model organisms, and an H-statistic of 1,352.19 with a p-value < 0.001 in pathogen proteomes. Nevertheless, we report significant differences depending on the organism studied and the taxonomy; for instance *Schizosaccharomyces pombe* shows a preference for taxonomic quasi-primes in Alpha helices and *Plasmodium falciparum* in Disordered secondary structures (Figure 9a-b). These findings showcase the structural diversity and potential functional significance of taxonomic quasi-prime peptides.

**Figure 9:**
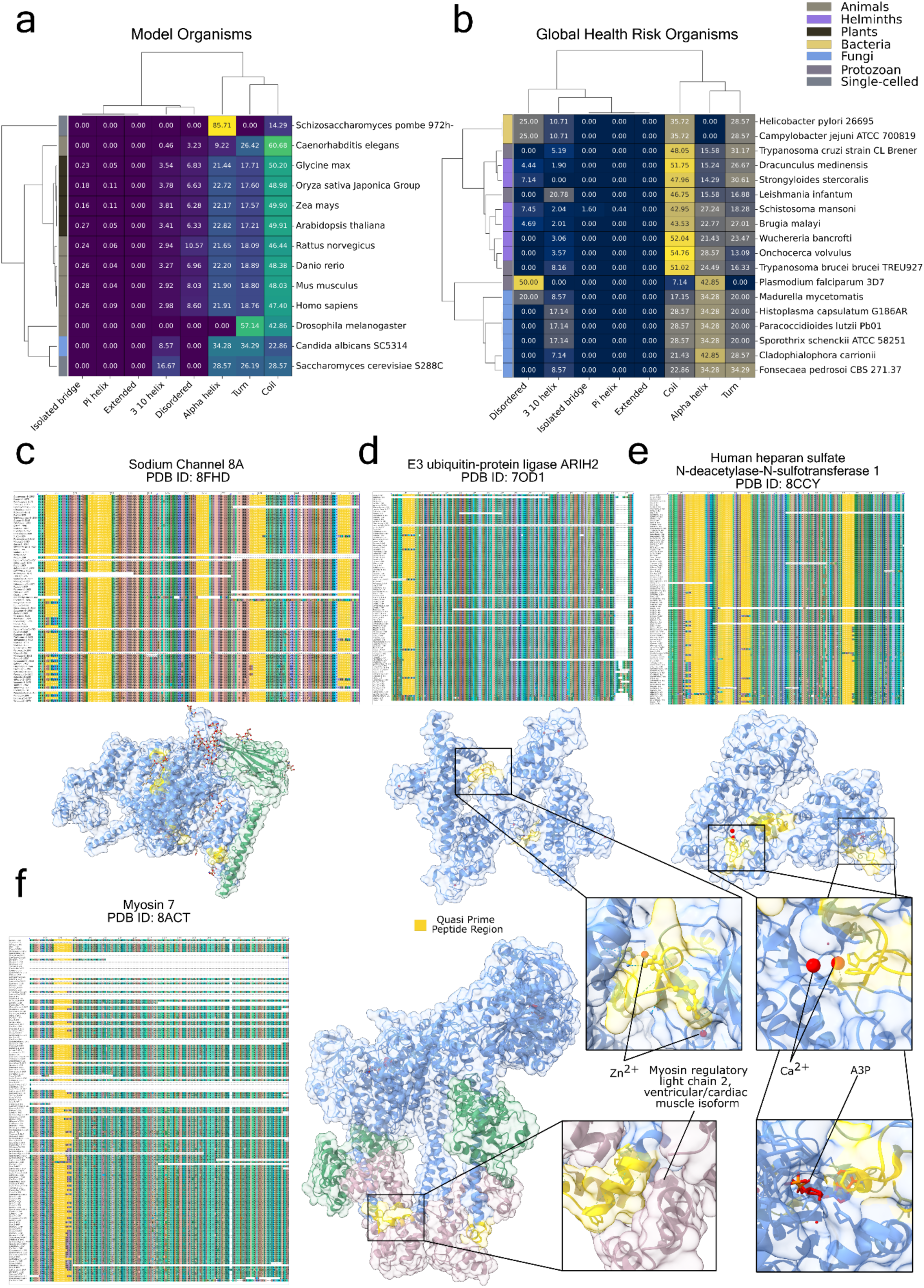
Structural profiling of -ε quasi-primes across evolutionary and pathogenic contexts. **a-b** Secondary structure profiling of taxonomic quasi-primes in model and global health risk organisms. **a** Clustered heatmap illustrating the secondary structure composition of taxonomic quasi-primes with ε exceeding 90% across selected model organisms. **b** Clustered heatmap of the secondary structure composition of taxonomic quasi-primes with ε values over 90% in selected global health risk organisms, encompassing bacteria, fungi, helminths, protozoans, and other single-celled pathogens. Rows (organisms) and columns (secondary structure types) have been hierarchically clustered based on Euclidean distance, employing Ward’s method to reveal patterns and structural similarities among species. Each heatmap cell displays the exact percentage of a specific secondary structure type within each organism. Multiple sequence alignment and structural representations of proteins containing taxonomic quasi-primes with ε-score greater than 90% in: **c** Human Sodium channel 8A (*SCN8A*), **d** Human E3 ubiquitin-protein ligase ARIH2 (*ARIH2*), **e** Human heparan sulfate N-deacetylase-N-sulfotransferase 1 (*NDST1*), **f** Human Myosin 7 (*MYH7*). Quasi-prime peptide region is displayed in yellow. Inset panels provide a detailed view of these regions, illustrating their specific locations within proteins, where they play a functional role. A3P: Adenosine-3’-5’-Diphosphate

### Multiple Sequence Alignment and Structural Insights into Chordata Proteins

Building on the aforementioned findings, we selected specific human proteins to investigate the functions of their corresponding quasi-primes in greater detail. We focused our study only on quasi-prime seven-mers with an ε-score greater than 90%, ensuring high confidence in functional relevance (**Supplementary Table 4**). To maintain structural accuracy, we analyzed only protein tertiary structures that were experimentally validated through high-resolution methods such as Cryo-EM or X-ray crystallography. Multiple Sequence Alignment (MSA) was conducted on corresponding protein orthologs across the Chordata species, allowing us to visualize the conservation of quasi-prime regions and to understand evolutionary patterns and functional conservation.

To examine the ion transport function, which is predominantly enriched in Eukaryota, we focused on Sodium Channel 8A (*SCN8A*) (Figure 9c), as the sodium channel α8 family exhibited notable enrichment in Chordata. For processes involving metal ion binding, we selected the E3 ubiquitin-protein ligase ARIH2 (*ARI2*) (Figure 9d), leveraging its association with the enriched TRIAD supradomain and its membership to the E3 ubiquitin ligase RBR family found enriched in the Chordata species, and the heparan sulfate N-deacetylase-N-sulfotransferase 1 (*NDST1*) (Figure 9e), chosen due to the enrichment of the heparan SO_4_ deacetylase domain. Lastly, to explore the role of quasi-primes in myosin-related functions and domains, we analyzed Myosin 7 (*MYH7*) (Figure 9f), providing further insight into the functional adaptations of these key proteins.

Twelve taxonomic quasi-primes, spread across various regions of the SCN8A protein, were identified (Figure 9c), suggesting a predominantly structural role rather than a functional one, as these regions do not actively interact with lipid substrates. In the ARI2 protein, three quasi-primes were found to interact directly with zinc ions, through coordinate covalent bonds at the Cys257, Cys260 and His265 sites, which are essential for the structural stability and catalytic activity of the enzyme (Figure 9d), since they are present in the IBR linker domain (Duda et al. 2013). Eleven taxonomic quasi-primes are identified in the NDST1 protein at regions related to calcium and adenosine-3’-5’-diphosphate (A3P). Particularly, the residues His389 and His393 both interact with the calcium ion through coordination covalent bonds, while Phe816 interacts with A3P through π-stacking interactions (Figure 9c). Quasi-primes present in both the sulfotransferase active site and the deacetylase active site, where they stabilize the structure enabling the correct placement of the N-acetyl-heparan sulfate (Mycroft-West et al., 2024). Additionally, five quasi-primes were identified in the MYH7 protein, interacting with the myosin regulatory light chain 2, ventricular/cardiac muscle isoform (MLC2v) (Figure 9d) through one hydrogen bond at the Trp827 site and electrostatic interactions at the positively charged Lys831. This interaction regulates motor function, stabilizes the myosin complex, and enhances calcium sensitivity, ensuring efficient cardiac muscle contraction (Rayment et al. 1993). These functions are critical for proper force generation and adaptability of the heart, with dysregulation linked to cardiomyopathies. These findings highlight the significance of quasi-prime detection in understanding protein interactions and their implications for healthcare.

### Pathogenic variants are enriched at human taxonomic quasi-prime loci

Finally, we examined if taxonomic quasi-prime loci are more likely to harbor pathogenic variants than surrounding sequences. We used human proteome-wide missense variant effect prediction maps (J. Cheng et al. 2023) and investigated if the subset of missense variants that are pathogenic are more likely to be found at taxonomic quasi-prime loci in humans. We find that the distribution of variants overlapping taxonomic quasi-primes is significantly sifted towards pathogenic effects (Kolmogorov-Smirnov test p-value<0.001, Cliff’s delta = 0.448) with taxonomic quasi-prime loci being 2.08-fold more likely to overlap pathogenic variants than expected (Figure 10). The increased pathogenicity at taxonomic quasi-prime loci, reflects their significance in traits and characteristics that are taxon-specific.

**Figure 10:**
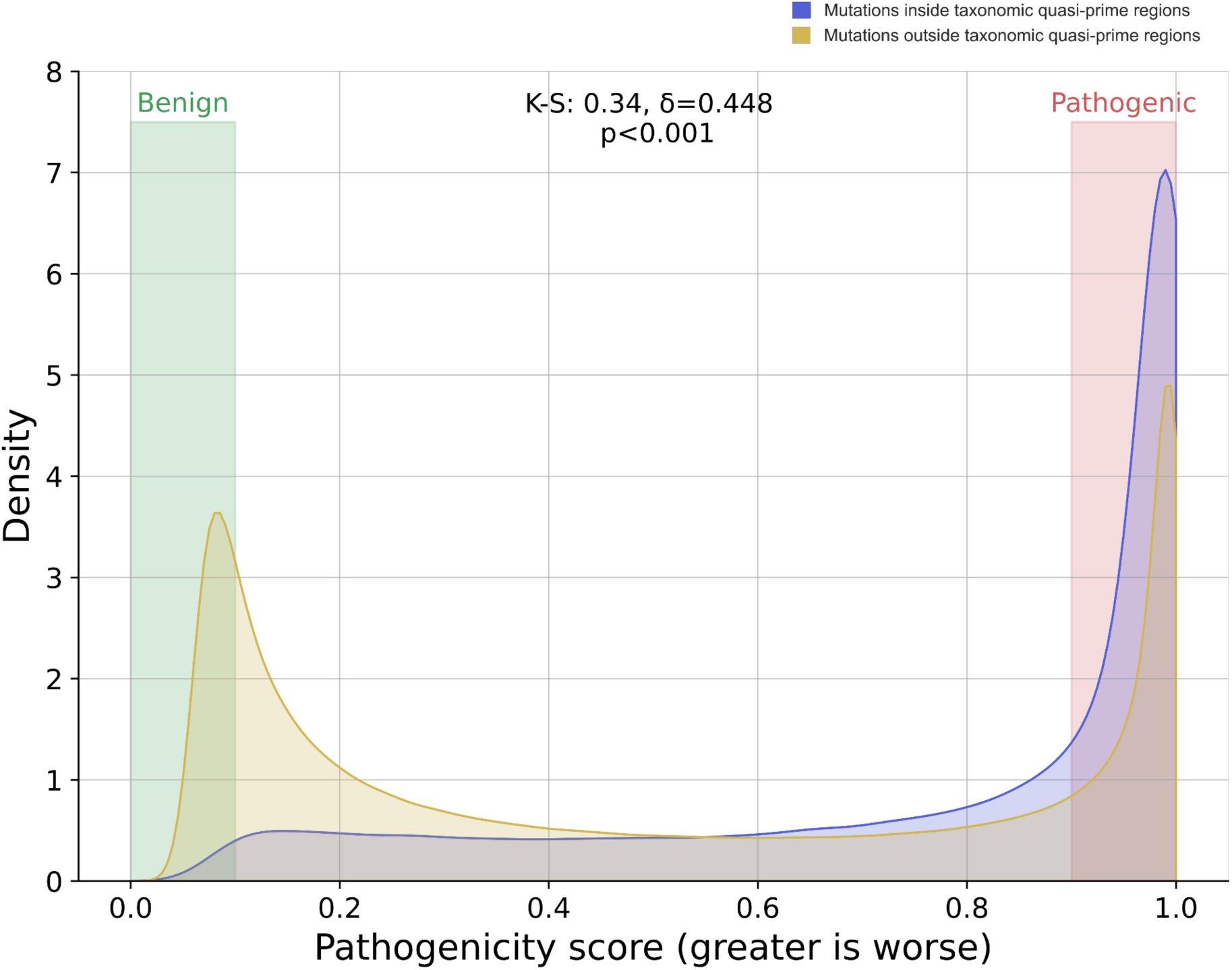
Missense mutations in highly conserved quasi-prime regions are more likely to be pathogenic. The Kernel Density Estimate (KDE) plot compares the distribution of AlphaMissense-predicted pathogenicity scores for missense mutations within human quasi-prime loci (for Chordata quasi-prime 7-mers with ε-scores above 90%) to those outside these loci within the same proteins. Highlighted ranges indicate highly benign mutations (pathogenicity scores < 0.1) and highly pathogenic mutations (pathogenicity scores > 0.9). Statistical metrics: Kolmogorov-Smirnov (K-S) statistic, Cliff’s delta (δ), and p-value (p).

## Discussion

The presence of short peptide k-mers that are highly specific to individual taxonomies has not been studied to date. Here, we provide evidence that based on the tens of thousands of available reference proteomes, such sequences can be systematically discovered across taxonomic levels. We identify peptides that exhibit remarkable taxonomic uniqueness at six- and seven-amino-acid lengths, offering insights into evolutionary divergence and lineage-specific adaptations. This study highlights taxonomic quasi-prime peptides as a novel approach for advancing our understanding of evolutionary processes and could be useful in developing innovative solutions in healthcare, agriculture, and environmental sustainability.

We observe large variations in the number and frequency of taxonomic quasi-primes when comparing different taxonomies. Protista display substantially lower ε_max_ scores when compared to other eukaryotic kingdoms, indicating that their characteristic quasi-prime peptides appear only in a small subset of the corresponding species. This can be attributed to the highly diverse polyphyletic nature of Protista, which encompasses both multicellular and unicellular organisms. Expanding on this, Protista exhibit a wide variety of environments, from marine and terrestrial ecosystems to parasitic niches, increasing the need to adopt different mechanisms for survivability (Burki, Sandin, and Jamy 2021). In bacteria and archaea, we find that taxonomic quasi-prime peptides are enriched in proteins associated with redox reactions. Eukaryotic phyla, such as Chordata and Streptophyta, displayed the largest number of enriched BPs which is in accordance with their larger proteome sizes and diverse developmental pathways.

The identification of taxonomic quasi-primes offers a novel way to study sequence divergence, speciation, and trait development. By identifying k-mer peptides unique to specific taxa, these sequences can serve as molecular markers of evolutionary processes and lineage-specific adaptations. Analyzing the proteins harboring quasi-prime peptides may reveal how unique traits emerge, such as specialized metabolic pathways or complex physiological processes. Future research could integrate phylogenetic analyses with quasi-prime peptide distributions to trace evolutionary histories and uncover the genetic basis of speciation events, shedding light on how sequence-level variations drive biological diversity and the evolution of novel traits.

Taxonomic quasi-prime peptides could have a wide range of applications in biological research, agriculture, and healthcare. Immunological methods based on antibodies can face limitations in sensitivity and specificity, particularly due to cross-reactivity (Wild 2013). By prioritizing antigens that harbor taxonomic quasi-prime peptides, the sensitivity and specificity of designed antibodies can be increased. Given their short length and unique taxonomic specificity, taxonomic quasi-prime peptides are well-suited for integration into mass-spectrometry workflows (Dupree et al. 2020; Birhanu 2023). Their inherent specificity can enhance the precision of proteomic analyses, enabling more efficient identification of proteins in complex biological samples (Li et al. 2023). In agriculture, taxonomic quasi-prime peptides can serve as highly-specific biomarkers for pathogen detection. Examination of quasi-prime peptides specific to microbial communities or ecological niches could also provide insights into microbiome dynamics and their role in ecosystem function. In healthcare, taxonomic quasi-prime peptides could facilitate the identification of pathogenic organisms in clinical samples, contributing to faster and more precise diagnostics for infectious diseases.

The number of available proteomes represents only a small subset of the species known. Therefore, future work is required to examine how these findings change as more reference proteomes of different organisms become available. Furthermore, the incorporation of protein isoforms and population variants could influence our conclusions and further exploration towards these directions is needed, particularly in eukaryotic proteomes, when such data become available across multiple organisms.

## Supporting information

Supplementary Table 6: Complete protein family names can be found at: families_short_names.csv

Supplementary Table 5: Complete protein domain names can be found at: domains_short_names.csv

## Code availability

The code for taxonomic quasi-prime and taxonomic nullomer extraction and subsequent analysis can be found at: https://github.com/Georgakopoulos-Soares-lab/taxonomic_quasi_primes

Raw taxonomic quasi-prime extraction data are available at: https://zenodo.org/records/14385095

## Acknowledgements

Research reported in this publication was supported by the National Institute of General Medical Sciences of the National Institutes of Health under Award Number R35GM155468. The content is solely the responsibility of the authors and does not necessarily represent the official views of the National Institutes of Health.

## Supplementary Material

**Supplementary Figure 1:**
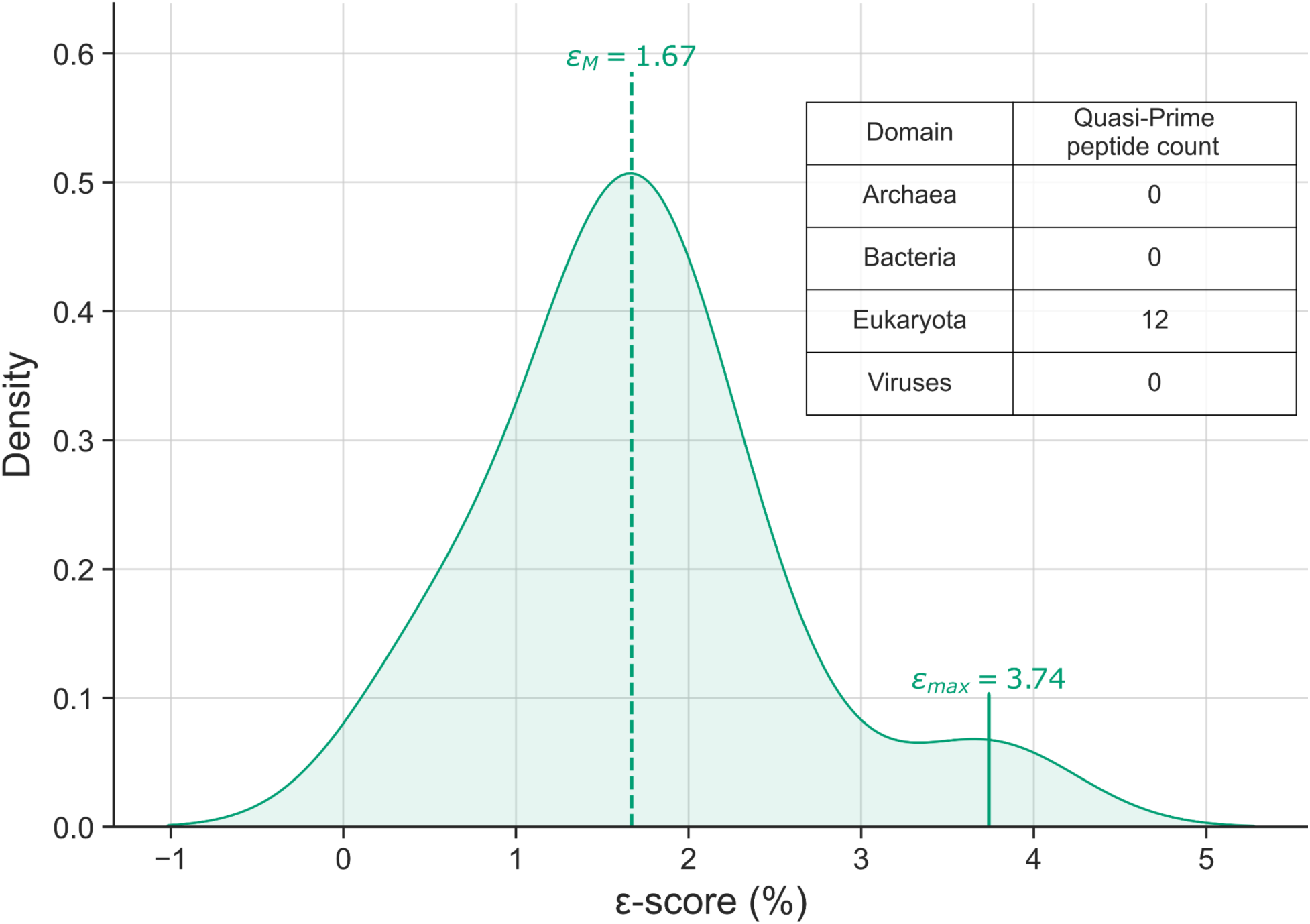
Kernel Density Estimate (KDE) plot illustrating the ε-score distribution of superkingdom quasi-prime 5-mers. The smoothing of the curve extending into negative values and exceeding εmax is likely due to the limited sample size of peptides, which affects the Gaussian kernel’s estimation fidelity.

**Supplementary Figure 2:**
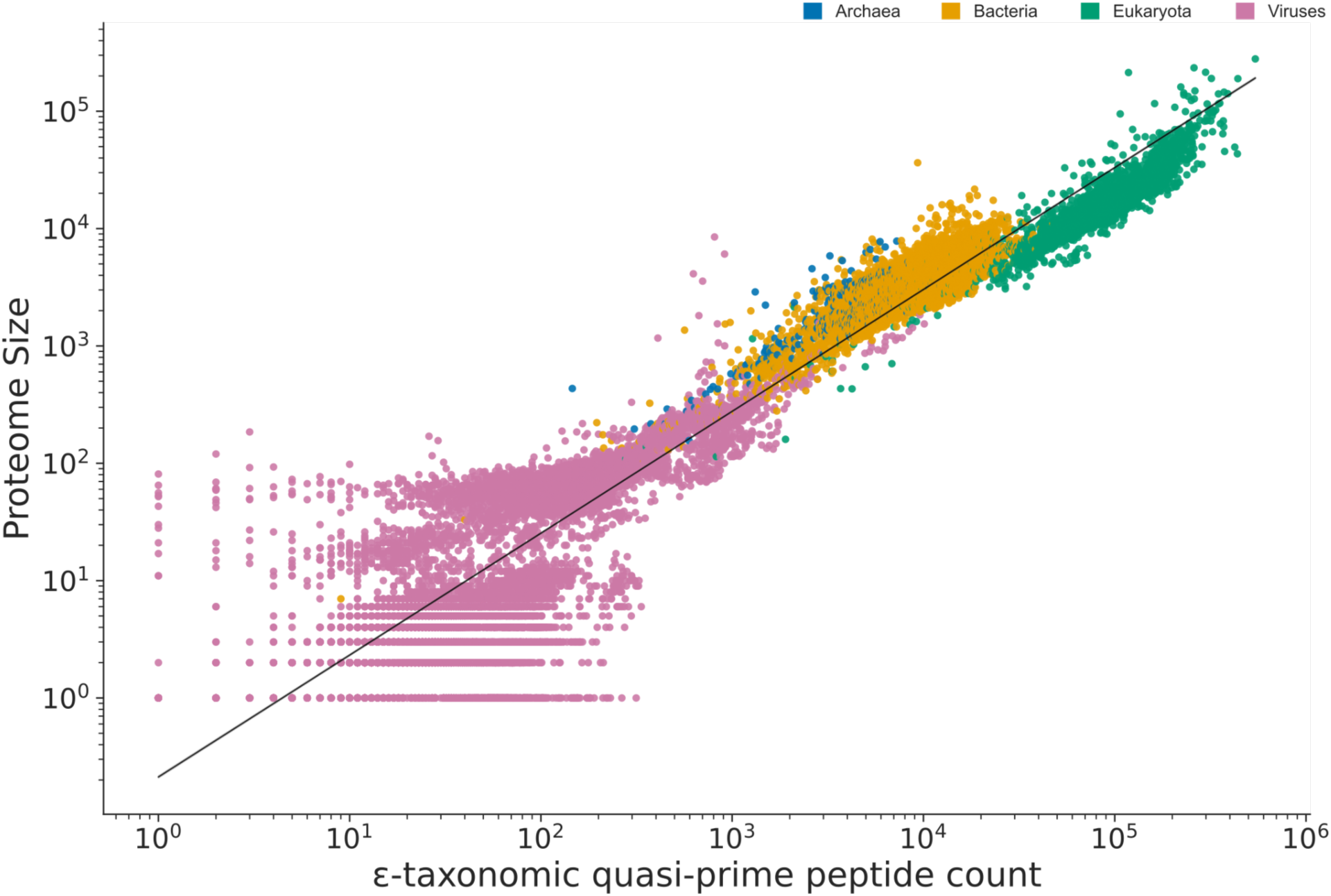
Scatterplot representing the number of unique taxonomic quasi-prime peptides (6mers and 7mers) observed in each reference proteome as a function of their proteome size.

**Supplementary Figure 3:**
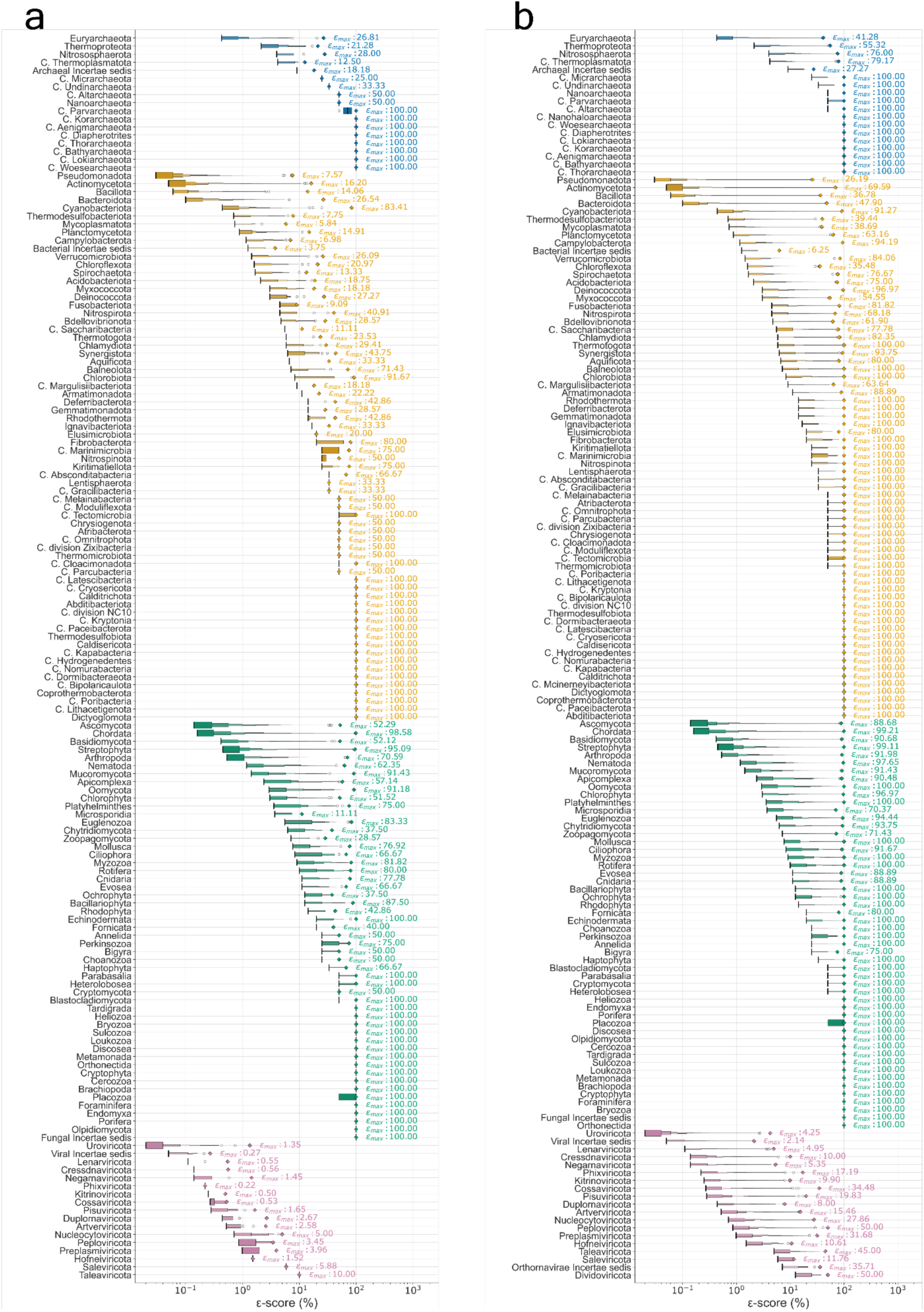
ε-score distribution of quasi-prime peptides across all phyla at the superkingdom level.

**Supplementary Table 1:**
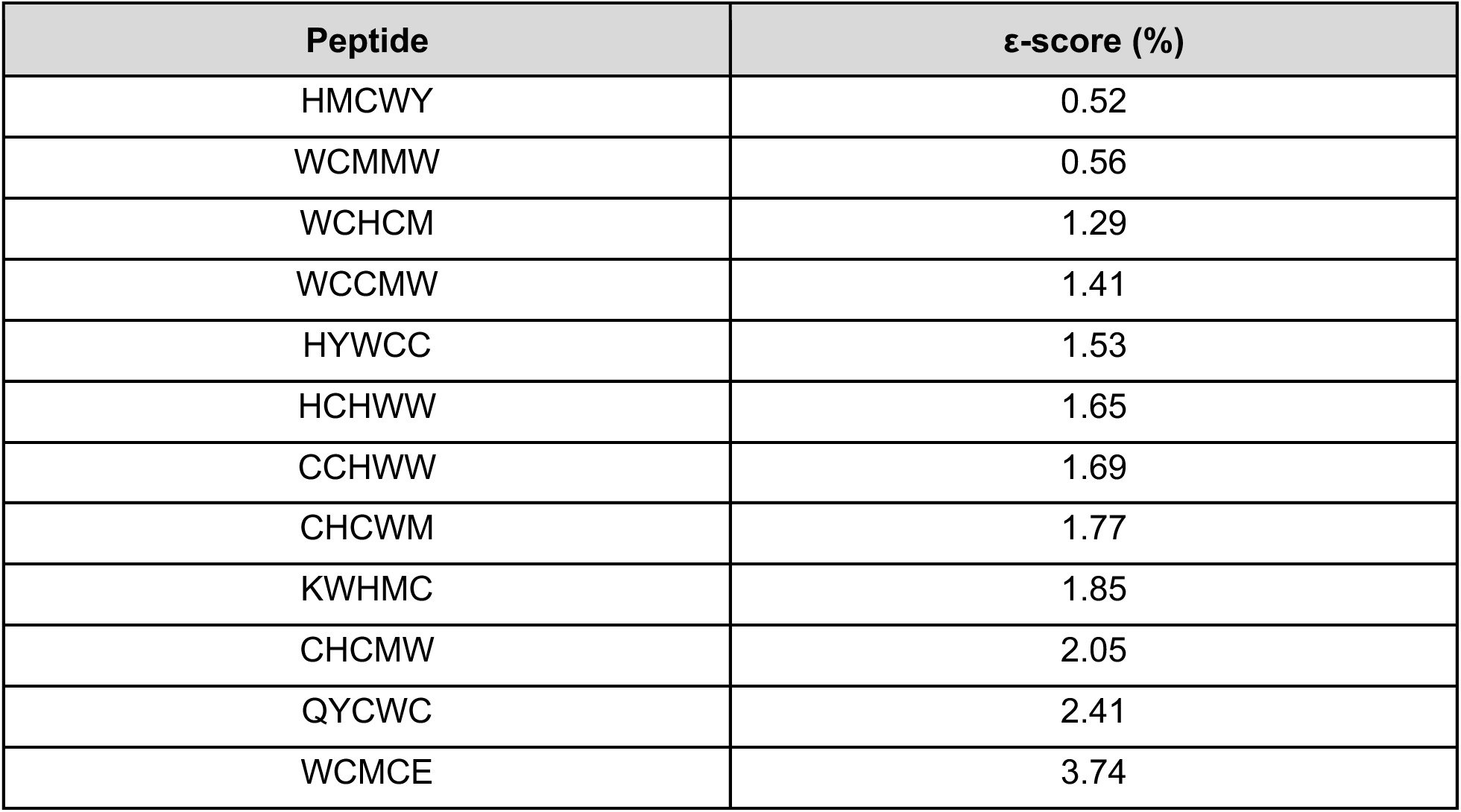
List of Eukaryota quasi-prime 5-mer sequences identified at the superkingdom level, along with their corresponding ε-scores.

| Peptide | ε-score (%) |
| --- | --- |
| HMCWY | 0.52 |
| WCMMW | 0.56 |
| WCHCM | 1.29 |
| WCCMW | 1.41 |
| HYWCC | 1.53 |
| HCHWW | 1.65 |
| CCHWW | 1.69 |
| CHCWM | 1.77 |
| KWHMC | 1.85 |
| CHCMW | 2.05 |
| QYCWC | 2.41 |
| WCMCE | 3.74 |

**Supplementary Table 2:** Quasi-prime peptides (six-mers and seven-mers) exhibiting the highest ε-score across Eukaryotic Kingdoms.

| Eukaryotic quasi-prime six-mers |  |  | Eukaryotic quasi-prime seven-mers |  |  |
| --- | --- | --- | --- | --- | --- |
| Kingdom | Peptide | $\epsilon_{\max}$ (%) | Kingdom | Peptide | $\epsilon_{\max}$ (%) |
| Fungi | FPKCYW | 34.58 | Fungi | DANQDNY | 92.50 |
| Metazoa | KWMMYW | 87.18 | Metazoa | CKGFFKR | 98.78 |
| Viridiplantae | YPCFMW | 82.88 | Viridiplantae | KSCRLRW | 98.05 |
| Protista | WHDCHC<br>WWEFYH | 17.03 | Protista | REENKWC | 37.91 |

Supplementary Table 3: εM values for 6mer and 7mer taxonomic quasi-primes across Phyla in each Superkingdom are available at:

phyla_summary.csv

**Supplementary Table 4:**
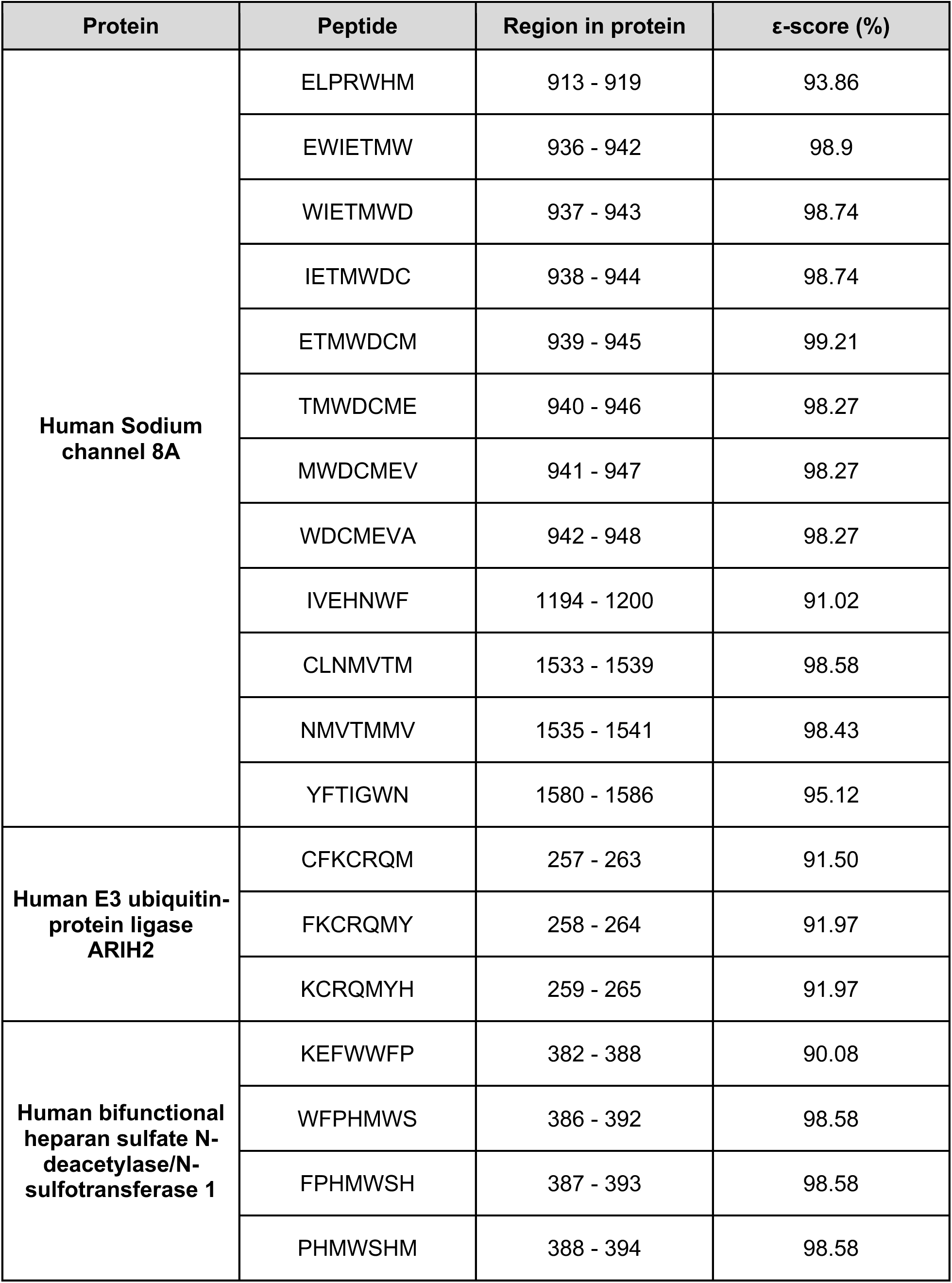
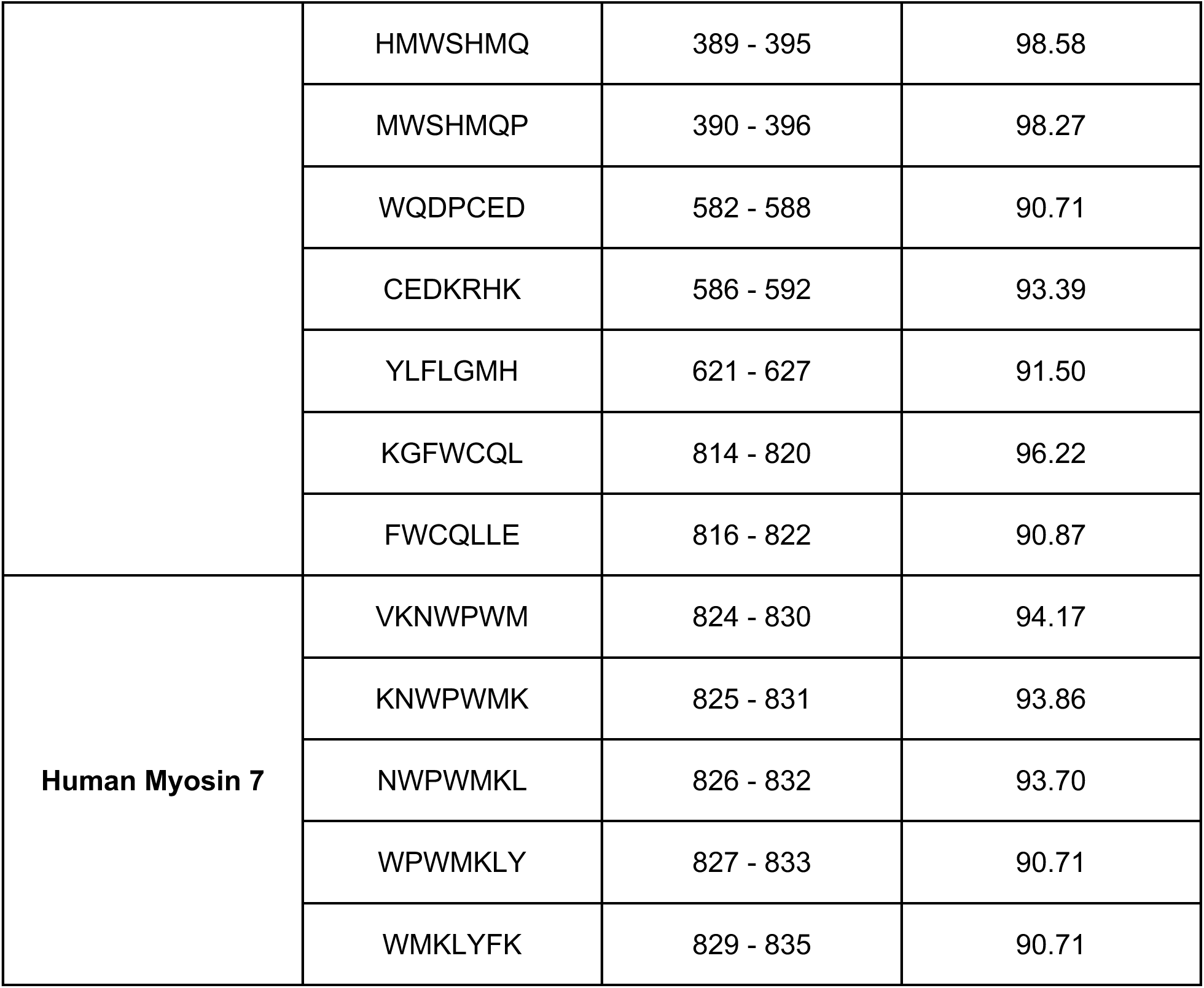
Quasi-prime 7mers identified in selected Chordata proteins.

**Supplementary Table 5: Complete protein domain names can be found at: domains_short_names.csv**

**Supplementary Table 6: Complete protein family names can be found at: families_short_names.csv**

